# Biocontrol efficacy of 2-pyrroldione-5-carboxylic acid (2Py-5CA), an antifungal bioactive macromolecule from the endophyte *Penicillium oxalicum*, against *Ramularia collo-cygni* in barley

**DOI:** 10.64898/2026.01.29.701693

**Authors:** Seema Rathore, Olga Lastovetsky, Sujit J. Karki, Junhao Xie, Angela Feechan

## Abstract

Global crop production suffers significant losses (23%) due to disease caused by fungal pathogens. Biocontrol agents may be sustainable alternatives to fungicides for disease management. Ramularia leaf spot (RLS), caused by *Ramularia collo-cygni (RCC),* is a widespread disease of barley (*Hordeum vulgare*). Two *Penicillium oxalicum* isolates identified from a collection of endophytes from wild barley, suppress development of *R. collo-cygni* and exhibit functional plant growth attributes *in vitro*. Analysis of the *P. oxalicum* culture filtrate revealed antifungal activity common to both isolates. An untargeted metabolomics (LC-MS/MS) approach uncovered the secondary metabolite profile of the *P. oxalicum* endophytes including the identification of 2Py-5CA as an antifungal. AntiSMASH-based genome mining identified a non-ribosomal peptide synthetase (NRPSs) biosynthetic gene cluster in *P. oxalicum* responsible for producing paraherquamide and ultimately the antifungal 2Py-5CA. *In vitro* studies demonstrated that 2Py-5CA can also suppress growth of the cereal pathogens; *Zymoseptoria tritici*, *Pyrenophora teres*, *Fusarium graminearum* and the horticultural pathogen, *Botrytis cinerea* disrupting the growth of pathogenic fungal hyphae and spore structure development. Finally, 2Py-5CA reduced RLS symptoms when sprayed on barley seedlings. Therefore 2Py-5CA has potential as a biofungicide to combat fungal crop pathogens.

**Graphic:** 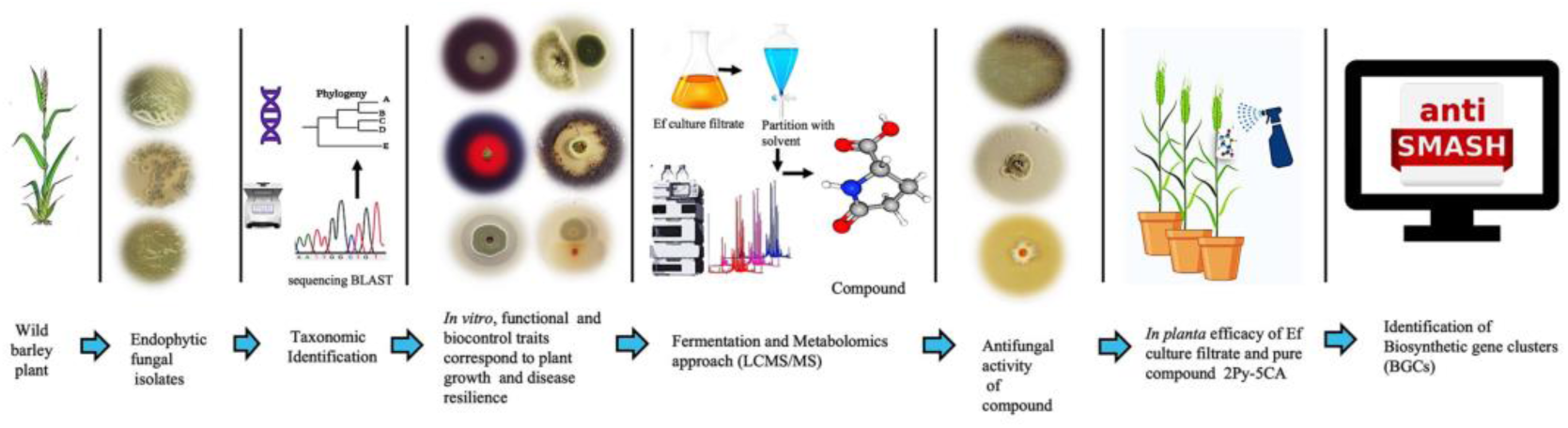

## 1. Introduction

Despite the use of chemical fungicides, global crop production suffers significant losses (23%) due to disease caused by fungal pathogens. This poses a major threat to food security [1] The United Nations Food and Agriculture Organization (FAO) estimates that growers must produce 70% more food by 2050 to meet the demands of the anticipated population increase [2]. At the same time, initiatives such as the EU Green Deal and the Farm to Fork strategy have set targets to reduce pesticide use by 50% by 2030 [3], [4]. Therefore, sustainable alternatives to chemical fungicides are essential to curb disease-associated crop losses and to ensure food security.

Barley (*Hordeum vulgare*) is the fourth-largest cereal crop globally, primarily used in feed and the brewing industry [5]. However, barley is susceptible to fungal pathogens that can cause substantial yield losses and impair grain quality [6]. *Ramularia collo-cygni (RCC),* the causative agent of Ramularia leaf spot (RLS) in barley, leads to reduced grain yield and quality by up to 70% across Europe and beyond [[7], [8]]. RLS symptoms include small brown lesions with chlorotic halos on leaves late in the season, reducing photosynthetic area [9], [10]. Chemical fungicides, such as quinone outside inhibitors (QoI), strobilurins, demethylation inhibitors (DMI), and succinate dehydrogenase inhibitors (SDHI), have become less effective against *R. collo-cygni* [11]. Moreover, the multi-site fungicide chlorothalonil was banned by the European Food Safety Authority (EFSA) in 2020 due to health and environmental safety concerns [12]. Given these challenges, there is a need to explore alternatives for crop disease control strategies [13]. A promising alternative to chemical fungicides is the use of biological control agents, including microbes such as fungal endophytes and their bioactive metabolites. Discovering natural endophytes as biocontrol agents provides a sustainable and eco-friendly alternative for managing plant disease, reducing reliance on synthetic fungicides, which are increasingly facing resistance development [14] Besides protecting plants from disease, fungal endophytes can also promote plant growth through various mechanisms [15], [16]. For example, *Trichoderma spp*. can be used in barley as a plant growth-promoting agent and for biocontrol against the pathogen *Pyrenophora teres* [17]. *Penicillium olsonii* ML37 and *Acremonium alternatum* ML38 can be implemented for integrated disease management in wheat against *Septoria tritici blotch* in the field [18].

Moreover, fungal endophytes have been shown to produce a wide range of secondary metabolites [19], [20]. Endophytic fungi are known to be a rich repository for bioactive molecules including terpenoids, flavonoids, alkaloids, and steroids, which possess diverse structures and biotechnological applications [21], [22]. For example, pyrrolidone is an important five-membered heterocyclic building block with one nitrogen atom which can lead to a wide variety of compounds with biological activities [23]. Pyrrolidione containing secondary metabolites include myrmicarin, lamellarin, ryanodine, makaluvamine M, all exhibiting potential biological activity [24], [26]. Various derivatives of pyrrolidione analogs have demonstrated a wide range of biological activities, including antifungal, antibacterial, anti-inflammatory, antioxidant, antiviral and anticancer effects[27], [32].

In this study, we report the isolation and characterization of an antifungal metabolite 2-Pyrrolidione-5-Carboxylic acid (2Py-5CA) isolated from *P. oxalicum* and evaluate its *in vitro* biocontrol potential against cereal pathogens. We used liquid chromatography with tandem mass spectrometry (LC-MS/MS) to identify 2Py-5CA from the culture filtrate of endophytic fungal isolates of *Penicillium oxalicum* (Ef20 and Ef30). The *Penicillium oxalicum* endophytes were obtained from a collection previously isolated from the wild barley *Hordeum secalinum and Hordeum murinum* [[33], [34]]. This antifungal metabolite 2Py-5CA has activity *in vitro* against a range of cereal pathogens including *R. collo-cygni, Zymoseptoria tritici*, *Pyrenophora teres* and *Fusarium graminearum* as well as the grey mould pathogen *Botrytis cinerea*, an important horticultural pathogen. *In planta* assays confirmed the application of 2Py-5CA effectively reduced RLS disease symptoms in barley. Furthermore, antiSMASH analysis identified the non-ribosomal peptide synthetase (NRPS) gene cluster, which is predicted to be responsible for the production of secondary metabolites. Taken together, these results provide a promising and potentially broad-spectrum biofungicide against fungal pathogens of cereal crops for sustainable plant disease management.

## 2. Materials and methods

### 2. 1. Endophytic Fungal Isolates

Using an *in vitro* confrontation assay, we screened 64 endophytes for their ability to inhibit the growth of *R. collo-cygni* from a previously isolated endophyte collection [33, 34]. Two isolates showed strong inhibition of *R. collo-cygni in vitro* designated as *Penicillium oxalicum* Ef20, *P. oxalicum* Ef30. Another isolate showed moderate inhibition, designated *P. virgatum* Ef65 (Fig S1; Table S1). Fungal isolates were grown on potato dextrose agar (PDA), incubated at 21 °C for 7 days and sub-cultured under the same conditions. A fresh-grown culture from the edge of each fungal isolate was grown in 100 mL of potato dextrose broth (PDB) and incubated at 21 °C on a rotary shaker for 4 days at 150 rpm. Stocks of endophytic fungal isolates were preserved in 50% v/v glycerol and stored at -80 °C.

### 2.2. Fungal Pathogen Isolates

Fungal pathogens used in this study: *Ramularia collo-cygni isolates (R. collo-cygni*) were obtained from Stephen Kildea, Teagasc, Crops Research Centre, Oak Park, Carlow. *Pyrenophora teres f. teres* isolate CP2189 and *Botrytis cinerea* Pers. (CBS120092) were obtained from the School of Natural Sciences and Trinity Centre for Biodiversity Research, Trinity College Dublin [35–36]. *Fusarium graminearum* wild-type strain GZ3639 were obtained from Prof. Fiona Doohan and the *Zymoseptoria tritici* isolate IPO323 was used [37–38]. These phytopathogens were used to assess the antifungal ability of endophytic fungal isolates and the 2Py-5CA metabolite isolated from *Penicillium oxalicum Ef20* and *Ef30.* The fungal pathogens were grown on PDA and incubated at 21 °C for *R. collo-cygni* or *Z. tritici* for 6 days and *F. graminearum, B. cinerea,* and *P. teres* were incubated for 4 days at 21 °C. Fungal mycelia were aseptically inoculated into potato dextrose broth (PDB) and incubated at 21 °C and 21 °C at 150 rpm for 4 days. Fungal pathogen stocks were maintained in glycerol 50% (v/v) and stored at -80 °C.

### 2.3. Molecular Identification of Fungal Endophytic Isolates

To identify endophytic fungi at the genus level, the internal transcribed spacer (ITS) regions of the rDNA were analyzed. Endophytic fungal isolates were cultured on PDB at 21 °C for 3 days. Genomic DNA was extracted using the DNeasy PowerSoil Pro Kit (QIAGEN) following the manufacturer’s protocol. The quality and quantity of the extracted DNA were assessed using a NanoDrop spectrophotometer. The ITS regions were amplified using the ITS1 (5’-TCCGTAGGTGAAC-3’) and ITS4 (5’-TCCTCCGCTTATTGATATGC-3’) primers. The PCR conditions were as follows: initial denaturation at 95 °C for 2 minutes, followed by 35 cycles of denaturation at 95 °C for 30 seconds, annealing at 60 °C for 30 seconds, extension at 72 °C for 4 minutes, and a final extension at 72 °C for 4 minutes. PCR products were visualized on a 0.8% agarose gel and further purified using the QIAquick Gel Extraction Kit (QIAGEN). The purified PCR product sequences were aligned using the BioEdit Sequence Alignment Editor [39]. The assembled consensus sequences were subjected to nucleotide BLAST for comparison with sequences that showed homology in the GenBank database. Multiple sequence alignments were performed using ClustalW, MEGA 11 software [40]. Phylogeny was reconstructed based on the ITS gene sequence using the Maximum Likelihood method with the PhyML programme under the General Time Reversible (GTR) substitution model [41]. The tree was rooted with sequences from *Aspergillus* .

### 2.4. *In vitro* confrontation assay against fungal pathogens

The antifungal activity of selected endophytic fungal isolates was evaluated against the barley phytopathogens *Ramularia collo-cygni*, *Pyrenophora teres* and *Fusarium graminearum*. To conduct the antifungal bioassays, the confrontation dual culture method was used on petri dishes containing PDA. Fungal plugs (5 mm) of the actively growing pathogens and agar plugs (2 mm) of the endophytic fungi were placed 3 cm apart from each other. All PDA Petri dishes were incubated at 21 °C under the dark and after 4 days the progression of fungal growth of *F. graminearum* and *P. teres* was measured [41]. The percentage inhibition of hyphal growth of the fungal pathogen (*P.teres, F. graminearum*) was determined by comparing the growth in the presence of endophytic fungi with that of the control using the formula; Inhibition (%) = Rc-Ri/ Rc × 100. Where Rc is the radial hyphal growth diameter of the fungal pathogen in the control plates (mm)- Ri is the radial hyphal growth of the fungal pathogen in the presence of endophytic fungi (mm) [42]. The experiment was repeated three times independently with three replicates each time. Culture of *R. collo-cygni* was grown for 4 days at 21 °C with shaking at 125 rpm. The culture was centrifuged at 4000 rpm for 5 minutes to collect fungal hyphae, the supernatant was discarded. The pellet was washed twice with double distilled water and resuspended in 10 mL of ddH_2_O. 100 µL of *R. collo-cygni* containing 4 × 10^6^ hyphae was spread evenly onto PDA plates and allowed to dry for 5-10 minutes. Four-day-old endophytic cultures were inoculated at the centre of the PDA plates pre-inoculated with *R. collo-cygni*. All the plates were incubated at 21 °C in the dark for 3-4 days. The diameter of the inhibition zone against *R. collo-cygni* colonies was measured in millimetres (mm). The zone of inhibition (ZOI) was calculated using the formula: *(Colony diameter of the endophytic fungi + clear zone diameter towards the RCC) **/** Colony diameter of endophytic fungi* [42]. The experiment was repeated three times independently with three replicates each time.

### 2.5. Determination of nutrient mobilisation

Nutrient mobilisation by the endophytic fungi was evaluated based on three parameters: phosphate solubilization, siderophore production, and ammonia production. These experiments were repeated three times independently, with each repetition consisting of three replicates.

#### 2.5.1. Phosphate solubilization activity

The phosphate solubilization ability of endophytic fungal isolates (Ef20, Ef30, and Ef65) was screened using Pikovskaya’s (PKV) agar [43]. Seven-day-old endophytic cultures were placed at the centre of Petri dishes containing PKV agar and incubated at 28°C. After 3 days the diameter of the halos around the fungal colonies was measured in millimetres (mm). The phosphate solubilization index was calculated using the formula: *Phosphate solubilization index = (Colony diameter +clear zone diameter) **/** colony diameter* [43].

#### 2.5.2. Ammonia production

The fungal isolates were grown in peptone water on a rotary incubator shaker at 150 rpm for 3 days at 21 °C. Following incubation, the culture was centrifuged, and 4-5 drops of Nessler’s reagent were added to the supernatant. A faint yellow colour indicated minimal ammonia production, while a deep yellow to brownish colour indicated maximal ammonia production [44].

#### 2.5.3. Siderophore production assay

Siderophore production was detected using the Chrome Azurol S (CAS) agar assay as described by Schwyn and Neilands [45]. Fungal isolates were precultured on PDA for 5-7 days at 28°C in the dark. Agar plugs (1 mm in diameter) were cut from the edges of fresh colonies using a sterile 10 µL tip and inoculated onto CAS agar medium. The plates were incubated in the dark at 28°C for 3 days. The appearance of an orange halo around the colonies indicated positive siderophore production. The diameter of the halos was measured using the same formula as for phosphate solubilization.

#### 2.5.4. Extracellular Enzymatic Potential

The extracellular enzymatic potential of endophytic fungal isolates was qualitatively estimated for cellulase, protease, lipase, and gelatinase production using plate assays. Fresh fungal cultures were grown on PDA, and 1 mm diameter agar plugs were cut using a 10 µL tip and placed onto specific solid media containing the substrates of interest. Protease production was assessed on skim milk agar (SMA) using the method described Ibrahim et al., [46]. Cellulase Production was tested on carboxymethyl cellulose (CMC) agar medium and Lipase Production was evaluated on peptone agar medium supplemented with 2% Tween 20. All plates were incubated at 28 °C for 3 days. At the experiment end cellulase production plates were flooded with iodine solution to visualize halo formation [46]. The appearance of a halo around the fungal colonies was considered a positive result for the production of protease, lipase, and cellulase.

#### 2.5.5. Indole acetic acid (IAA) production

Indole acetic acid (IAA) production was assessed using PDB supplemented with L-tryptophan. A 100 mL volume of PDB, containing 0.5% L-tryptophan, was inoculated with endophytic fungal isolates and incubated on a rotary shaker at 150 rpm and 21°C for 4 days. After incubation, the culture broth was centrifuged at 4000 rpm for 5 minutes, and the supernatant was collected for IAA analysis. To detect IAA production, 1 mL of the supernatant was mixed with 2 mL of Salkowski reagent (prepared by combining 1 mL of 0.5 M FeCl₃ with 50 mL of 35% perchloric acid). The mixture was incubated in the dark for 30 minutes. The development of pink to the brick-red colour indicated the presence of IAA. The absorbance of the samples was measured at 530 nm using a spectrophotometer. Uninoculated growth medium served as a control. The concentration of auxin produced by the endophytic fungal isolates was estimated by extrapolating the absorbance values against a standard curve of IAA [47].

### 2.6. Light microscopy observation of 2Py-5CA antifungal compound on pathogenic fungi

The Interactions between endophytic fungal isolates and fungal pathogens, hyphal morphology were observed using light microscopy (Leica DM5500B microsystem). To investigate the interactions between endophytic fungal isolates and fungal pathogens, samples were collected from the inhibition zone periphery, placed onto sterile glass coverslips, hyphae and conidia stained with lactophenol cotton blue (Sigma-Aldrich).

### 2.7. Antifungal activity of *P. oxalicum* Ef30 culture filtrate

*P. oxalicum* Ef30, Ef20 and Ef65 were cultured in PDB on an incubator shaker at 150 rpm, 21°C for 7 days. After incubation, the culture was centrifuged at 4000 rpm for 10 minutes at 21°C and the mycelia discarded. The culture filtrate supernatant was collected, filtered aseptically through a 0.45 µm pore size membrane filter and added to fresh PDA media in 6-well plates at a final concentration of 16% (v/v). Each culture filtrate was tested in triplicate. Each well in the 6-well plate was inoculated with *R. collo-cygni* and its growth assessed relative to control (PDA with no added culture filtrate) after 7 days at 21 °C under dark conditions.

### 2.8. *Ramularia Leaf Spot (RLS)* disease assay in barley

Barley (*Hordeum vulgare*) cultivar RGT Planet seeds were sown directly into plastic pots with John Innes Compost No.2 potting soil from Westland Horticulture, UK. The growth chamber was set to a 16/8 h light/dark photoperiod, with relative humidity maintained at 80-100% and a temperature of 20°C. To assess the biocontrol efficacy of the *P. oxalicum* (Ef30) culture filtrate and pure compound 2-Pyrrolidone-5-carboxylic acid (2Py-5CA) against *Ramularia collo-cygni* (RCC) *in planta*, foliar spray application was employed.

Fourteen-day-old barley seedlings were inoculated by spraying the second leaf (GS12) with 5 mL of a *R. collo-cygni* spore suspension. The fungal hyphae of *R. collo-cygni* cultured in potato dextrose broth (PDB) were centrifuged (4000 rpm, 21°C, 5 minutes), washed, filtered using J-cloth, resuspended in distilled water (ddH2O), and sonicated. *R. collo-cygni* suspension (5 mL) was applied to each plant using handheld spray bottles until runoff. Immediately after inoculation, plants were covered with transparent plastic bags to maintain high relative humidity (80-100%) for 48 hours. After 48 hours, the bags were removed followed by treatment with the *P. oxalicum* (Ef30) culture filtrate or 2Py-5CA (10 µg/mL, 20 µg/mL, 40 µg/mL, and 80 µg/mL). Plants were sprayed with Tween 20® solution (1µL/mL) or with 0.5% DMSO respectively as a control. Disease scoring was performed on the second leaf by recording the relative area covered by disease symptoms. Scoring was conducted at multiple time points post-infection (8, 10, 13, 15, 17, and 21 days post-inoculation (dpi)). Disease severity (DS) levels was assessed using a visual 5-level scale: 0 = no visible symptoms; 1 = <10% of leaf area infected (necrotic lesions and chlorosis present); 2 = 10-25% infected area; 3 = 25-50% infected area; 4 = 50-75% infected area; and 5 = 75-100% infected area [48] Three independent experiments were conducted, each with two replicates per treatment. Each replicate included at least 10 leaves from 10 individual plants.

### 2.9. Discovery of Antifungal Metabolites in Culture Filtrate of Endophytic Fungi *Penicillium oxalicum* Ef30

#### 2.9.1. Metabolite extraction from *P. oxalicum* Ef30

For metabolite extraction, *P. oxalicum* Ef30 was inoculated into PDB and incubated for 7 days at 150 rpm and 21°C in a rotary shaker for the production of secondary metabolites. After incubation, the endophytic fungal culture was centrifuged at 4000 rpm for 10 minutes at 21°C and the fungal mycelium mat, contained within a cellophane layer, was discarded. The culture filtrate was extracted with an equal volume of ethyl acetate (1:1). This process resulted in two layers: the upper organic phase (ethyl acetate) and the lower aqueous phase. Each phase was collected separately into individual falcon tubes and lyophilized. Following lyophilization, the ethyl acetate phase was resuspended in 2 mL of methanol, and the aqueous phase was resuspended in 2 mL of sterile deionised water.

#### 2.9.2 Antifungal disc diffusion assays

The antifungal effectiveness of the ethyl acetate and aqueous metabolite fractions were assessed using a disc diffusion assay. The *R. collo-cygni* culture was grown on a rotatory shaker for 7 days at 21°C, 150 rpm in PDB. 100 µL of *R. collo-cygni* inoculum was spread evenly with a spreader on PDA and allowed to dry for 5 minutes. A sterile paper disc was placed in the centre of the PDA plate. For the evaluation of antifungal activity Ethyl acetate and aqueous extract were dissolved in methanol. Both ethyl acetate and water extract (50 µL) were applied to the paper disc, and an equal volume of methanol (50 µL) was applied. The plates were assessed for any clearing of *R. collo-cygni* growth around the disc. Each assay was repeated three times independently with three plates per replication.

### 2.10. Metabolomics (LC-MS/MS) based identification of bioactive compounds present in the aqueous extract

Extracts from the culture filtrates of *P. oxalicum* Ef20, *P. oxalicum* Ef30, *P. virgatum* Ef65 and *Penicillium verruculosus* Ef39 (as negative control) were dissolved in methanol, centrifuged at 1800 g for 5 mins, and filtered through a 0.22-μm filter. Liquid chromatography (LC/MS) and Electrospray ionisation (ESI) mass spectra analysis were performed using an Agilent 6545 QTOF LC/MS time-of-flight mass spectrometer, which consists of a 1290 Infinity II LC system and an Agilent Jetstream Electrospray Ionization (ESI) source coupled to a 6545 QTOF mass spectrometer. Liquid chromatography was performed on a Zorbax eclipse plus C18 column (2.1 x 50 mm, 1.8 µm) coupled to a Zorbax eclipse plus C18 guard column (2.1 x 5 mm, 1.8 µm). The column temperature was set at 30 °C, and the injection volume was 5 μL. The mobile phase A was water and B was 80% acetonitrile, both with 0.1% formic acid. The flow rate was 0.4 ml/min, and gradient conditions were set as follows: 1% B (0–1.5 min), 11% B (1.5–9 min), 25% B (9–15 min), 50% B (15–18 min), 99% B (18–18.05min), 99% B (18.05–21 min), 1% B (21–21.05 min) and 1% B (21.05–23 min). The MS parameters used for the analysis: drying gas temperature, 325 °C; drying gas flow rate, 10 l/min; sheath gas temperature, 350 °C; sheath gas flow rate,11 l/min; nebulizer pressure, 45 gauge pressure (pounds per square inch); capillary voltage, 3500 V; nozzle voltage, 1000 V; fragmentor voltage, 100 V; skimmer, 45 V. The mass-to-charge ratio (m/z) range was between 50 and 1600. Samples were run in both positive and negative ionization modes. MSI (mass spectrometry imaging) data was generated by Agilent Mass Hunter qualitative analysis software [49].

### 2.11. Metabolomics data processing

Data were acquired using MassHunter acquisition B.08.00 software (B.08.00.8058.3 Sp1Agilent Technologies) and were further processed in MassHunter qualitative analysis software (B.07.00 Sp2 Agilent Technologies). Molecular features were extracted using the molecular feature extractor (MFE) algorithm, and a list of features was generated with retention time, m/z, adducts or isotopes of compounds, signal intensity and accurate mass. MS/MS compound identification efforts included the fragment matches with the Human Metabolome Database (HMDB) and METLIN. Furthermore, the first 18 interesting features with high variable importance of projection (VIP) scores were generated. The exported data from Mass Profiler were analysed in SIMCA-P software (version 13.0.3; Umetrics) to calculate the retention time, m/z value, molecular mass and generate a list of compounds [50]. Compounds were compared between the endophytic fungi Ef20, Ef30 which inhibit *R. collo-cygni* growth, Ef65 which moderately inhibits *R. collo-cygni* and the negative control Ef39. Eighteen shared compounds present in the aqueous fraction of Ef20 and Ef30 only using LC-MS/MS were identified (Table S2). Five of these (2-Pyrroldione-5-carboxylic acid; 3-Methyl-2-oxovaleric acid; L-Cyclo(leucylprolyl); Mevalolactone; Pyridoxine), which are commercially available (Table S3) were tested for *R. collo-cygni* inhibition *in vitro*. The standards for 2-pyrroldione-5-carboxylic acid (2Py-5CA), 3-methyl-2-oxovaleric acid (3-Methyl-2-OVA), mevalonolactone, and pyridoxine were purchased from Sigma Aldrich, while L,L-cyclopropyl was obtained from Biosynth Ltd. Methanol and ethyl acetate of analytical grade (Merck). For LC-MS analysis, LC/MS grade acetonitrile and HPLC grade formic acid served as the mobile phase. Among the tested compounds , only 2Py-5CA demonstrated strong inhibitory activity against *R. collo-cygni*., A reference standard of 2Py-5CAwas used for comparison of the chromatographic peaks n the aqueous extract. Comparison was based on retention time (RT), mass-to-charge ratio (m/z), adduct formation, and isotopic pattern. Further confirmation was obtained through MS/MS fragmentation analysis with fragment spectra cross-referenced against the Human Metabolome database (HMDB) and METLIN [51].

### 2.12. Inhibition of fungal pathogen growth by 2Py-5C

A stock solution of 2Py-5CA was prepared by dissolving the 2Py-5CA in 0.5% DMSO. From this stock, a series of working concentrations (10, 20, 40, 80 and 100 µg/mL) were subsequently prepared for experimental use. 1 mL of each concentration of 2Py-5C was added to PDA to assess the antifungal activity against *R. collo-cygni, P. teres, F. graminearum, Z.tritici* and *B. cinerea* growth. The cultures of the pathogenic fungi *R. collo-cygni* and *Z. tritici* were grown for 4 days at 21 °C with shaking at 150 rpm. At the end the cultures were centrifuged at 4000 rpm for 5 minutes to collect the fungal hyphae and the supernatant was discarded. In the case of *R. collo-cygni* the fungal hyphae and *Z. tritici* blastospores were then washed twice with double-distilled water and subsequently resuspended in 10ml of ddH_2_O for experimentation. 100 µL of *R. collo-cygni* 4 × 10^6^ hyphae/mL and 5 × 10^6^ blastospores/mL suspension of *Z. tritici* were spotted onto a PDA plate and spread evenly. The plates were allowed to stand for 10-15 minutes. For other fungal pathogens, a mycelial plug (5 mm), from the perimeter of a six-day-old actively expanding colony of *P. teres*, *F. graminearum* and *B. cinerea* was placed at the centre of the PDA plate. The culture medium (PDA) with DMSO 0.5% was used as a control. Plates were incubated at 21 °C for 5-7 days for *R. collo-cygni* and *Z. tritici* growth and *P. teres*, *F. graminearum* and *B. cinerea* plates were incubated at 21 °C for 3-4 days in the dark. The growth of *R. colo-cygni* and *Z. tritici* colonies was observed to evaluate the different compound dosages using ImageJ. The surface area covered by *R.collo-cygni* and *Z.tritici* colonies treated with different concentrations of 2Py-5CA was quantified using ImageJ software (https://imagej.net/ij/). Photographs of *R. collo-cygni* and *Z. tritici* agar plates treated with 2Py-5CA were converted to 8-bit grayscale, and a uniform threshold was applied to distinguish colonies from background. Thresholder images were converted to binary (black for colonies, white for the background) and were excluded using a minimum particle size filter. The total surface area occupied by colonies was then measured using the “Analyse Particles” function after spatial calibration and results were expressed as colony-covered area (mm²) per plate. The total colony-covered surface area per plate was measured, averaged across triplicate plates, and exported to GraphPad Prism for statistical analysis.

For monitoring the percent radial growth of fungal pathogen hyphae *P. teres*, *F. grameniarum* and *B*. *cinerea* on plates with 2Py-5CA (10, 20, 40, 80, and 100 µg/mL) and control DMSO (0.5%) the percentage radial growth inhibition (PRGI%) was recorded. The experiment was repeated three times independently with three replications. The PRGI% was calculated as:

Percent radial growth inhibition of the hyphal diameter (PRGI%) = [(Rc -Ri)/ Rc] × 100

Where Rc is the radial hyphal growth of the fungal pathogen in the control plates (mm) - Ri is the radial hyphal growth of the fungal pathogen on2Py-5CA treated plates (mm). The results were analyzed by analysis of variance (ANOVA) using Tukey’s test.

### 2.13. Analysis of secondary metabolite biosynthetic gene clusters

For Whole Genome Sequence analysis, de novo assembly of sequence data was performed using Geneious Prime v2025.X (Biomatters Ltd., New Zealand) Cleaned FASTA-formatted read files (.giz) of *P. oxalicum* Ef20, *P. oxalicum* Ef30, and *P. verrculosus* Ef39 (used as a negative control which did not inhibit the *RCC in vitro*) genome sequences were imported into Geneious Prime (version 2025.0.3) for *de novo* assembly using integrated plugin tools. Sequences were pre-processed by trimming low-quality ends (Q-score threshold ≥30), removing adapter sequences using the BBMap adapter removal plugin, and filtering out reads shorter than 200 bp [52–53]. To investigate secondary biosynthetic potential of the endophytes, the antiSMASH (v7.0.1) pipeline was utilized with default relaxed parameters to identify biosynthetic gene clusters (BGCs) within the genomes of *P. oxalicum* Ef20, *P. oxalicum* Ef30, and *P. verruculosus* Ef39. The “Known clusterBlast” was used to identify homologous clusters across the analyzed genomes. Additional annotations were conducted using PFAM (protein families database) and the Minimum Information about a Biosynthetic Gene cluster (MIBiG) repository, which provided comprehensive metadata on BGCs and their associated products [54]. Furthermore, specific analyses of non-ribosomal peptide synthetases (NRPSs) and polyketide synthetases (PKSs) were carried out to predict the structural and functional diversity of the biosynthetic compounds encoded within these clusters.

### 2.14. Statistical analysis

All statistical analysis was performed using GraphPad Prism version 8.00 software (GraphPad Software, San Diego, CA, USA; https://www.graphpad.com/) to analyze the experimental data and create graphs. Data are expressed as mean ± SD. Statistical significance was calculated using a one-way analysis of variance, followed by Tukey’s multiple tests. *P-*value of < 0.05 was considered statistically significant.

## 3. Results

### 3.1. Endophytic Penicillium oxalicum Inhibits Ramularia collo cygni (RCC) growth

We screened 64 fungal endophytes previously collected from an endophytic library from different tissues of wild barley [33–34]. Ten isolates exhibited strong inhibition against *R. collo-cygni* (including Ef20 and Ef30), twenty-two isolates showed moderate inhibition (including Ef65), and twenty-three had no inhibitory effect (including Ef39). (Fig. S1) (Table S1). Furthermore, the percentage inhibitory effect was examined, by measuring the surface area covered by *R. collo-cygni* colonies in the presence of endophytic fungi, using ImageJ.

Based on this inhibition, we selected two of the strongest candidates (designated Ef20 and Ef30) which showed a significant inhibitory effect against *RCC*, while one candidate displayed a moderate effect Ef65 and a weak inhibitory candidate Ef 39 was used as a negative control (Fig. S1B). The percent surface area covered by *R. collo cygni* in the presence of the endophytic fungi Ef20 was 19.17% ± 1.00; Ef30 17.18% ± 0.74 ; Ef65 49.2% ± 2.11 and Ef39 61.57% ± 2.07 of the *R. collo cygni* only control (Fig. S1C).

To identify these endophytic fungi, colony morphology and conidia were examined (Figure S2). Molecular identification of Ef20, Ef30 and Ef65 was conducted via sequencing with ribosomal internal transcribed spacer (ITS) universal primers ITS1 and ITS4. Sequences retrieved from NCBI GenBank were aligned followed by the Neighbour-joining method to construct phylogenetic tree topologies. This phylogenetic analysis indicates Ef20 and Ef30 are *Penicillium oxalicum,* while Ef65 shares similarity with *P. virgatum* (Figure 1).

**Fig. 1.**
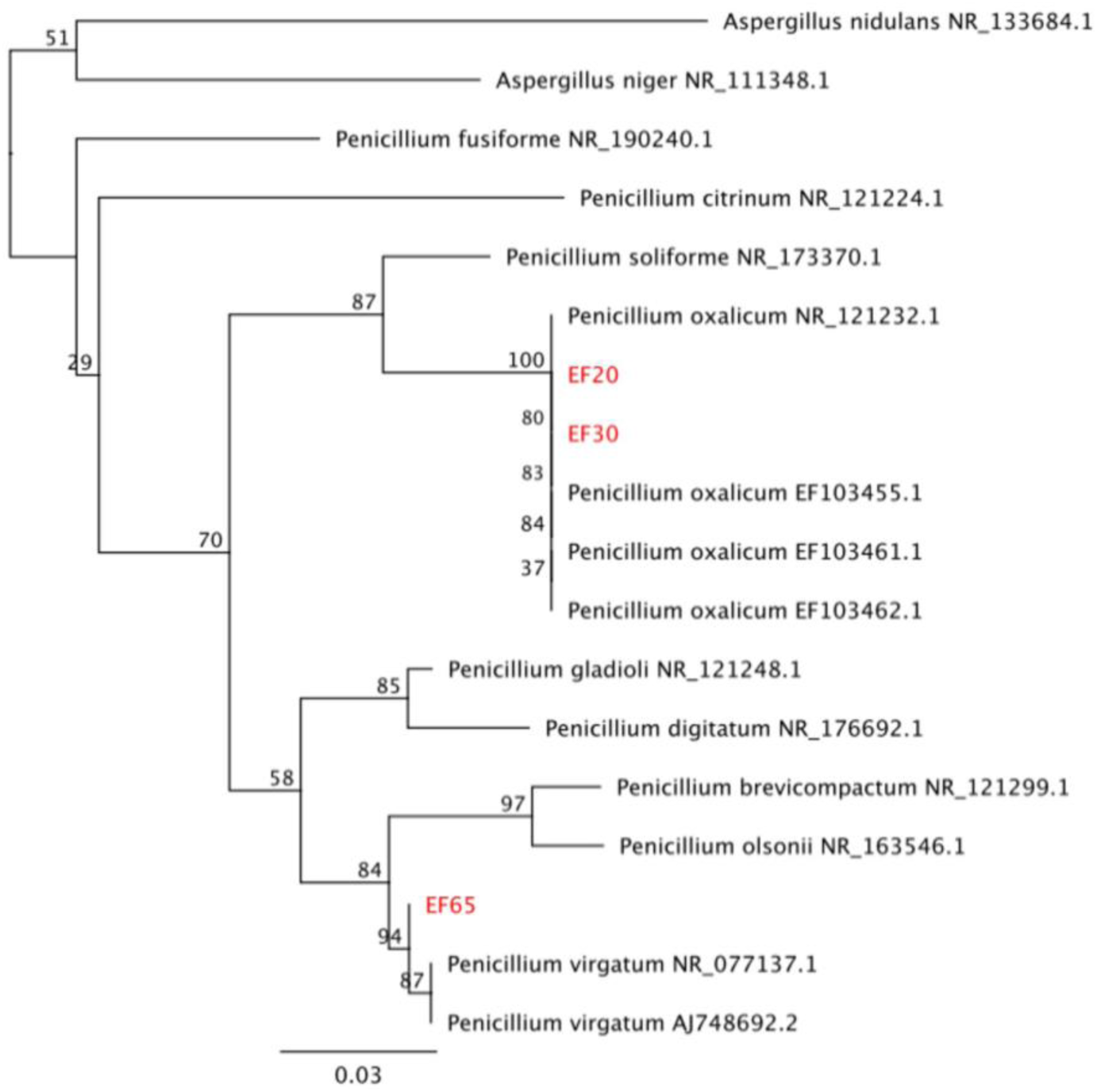
Phylogenetic placement of endophytes EF20, EF30 and EF65 in relation to known *Penicillium* spp. Phylogeny was reconstructed based on the ITS gene sequence using the Maximum Likelihood method with the PhyML programme (1) under the GTR substitution model. Maximum Likelihood bootstrap support values are displayed above branches. Values > 70% are considered significant. The tree was rooted with sequences from *Aspergillus* spp.

### 3.2. The Endophytic Fungi *Penicillium oxalicum* Possess Plant Growth-Promoting (PGP) Traits

To assess the potential of the fungal endophytes Ef20, Ef30 and Ef65 for the promotion of plant growth, the nutrient mobilization parameters: phosphate solubilization, siderophore production, and ammonia production were assayed. *P. oxalicum* Ef30 (clear zone of 34.67 ± 0.59 mm) and *P. oxalicum* Ef20 (28.33 ± 0.58 mm) both had significantly higher phosphate solubilisation compared to *P. virgatum* Ef65 (11.67 ± 0.62 mm) (Fig. 2, Table 1). IAA production was also significantly higher in *P. oxalicum* Ef30 (104 ± 2.54 µg/mL) and *P. oxalicum* Ef20 (103.3 ± 2.69 µg/mL) compared to moderate IAA production by *P. virgatum* Ef65 (31 ± 1.01µg/mL) (Table 1). Ammonia production was evaluated based on the ability to change the colour of the inoculation peptone growth medium by adding Nessler’s reagent. All the *Penicillium sp.* showed ammonia production changing the peptone growth medium colour from yellow to brown (Table 1). To assess the potential to assist plant iron uptake, siderophore production was evaluated using CAS-blue agar. The diameter of the orange clear zone observed for *P. oxalicum* Ef30 (35.0 ± 2.64 mm) and *P. oxalicum* Ef20 (30.7 ± 1.15 mm ) was significantly higher than for *P. virgatum* (17.33 ± 0.57 mm) (Fig. 2B, Table 1). Endophytic fungal isolates were qualitatively examined for extracellular enzyme synthesis, including cellulase, protease and lipase. Both *P. oxalicum* Ef20 and *P. oxalicum* Ef30 had cellulase, protease and lipase activity. Whereas, *P. virgatum* Ef65 had low cellulase and protease activity only (Table 1).

**Fig. 2.**
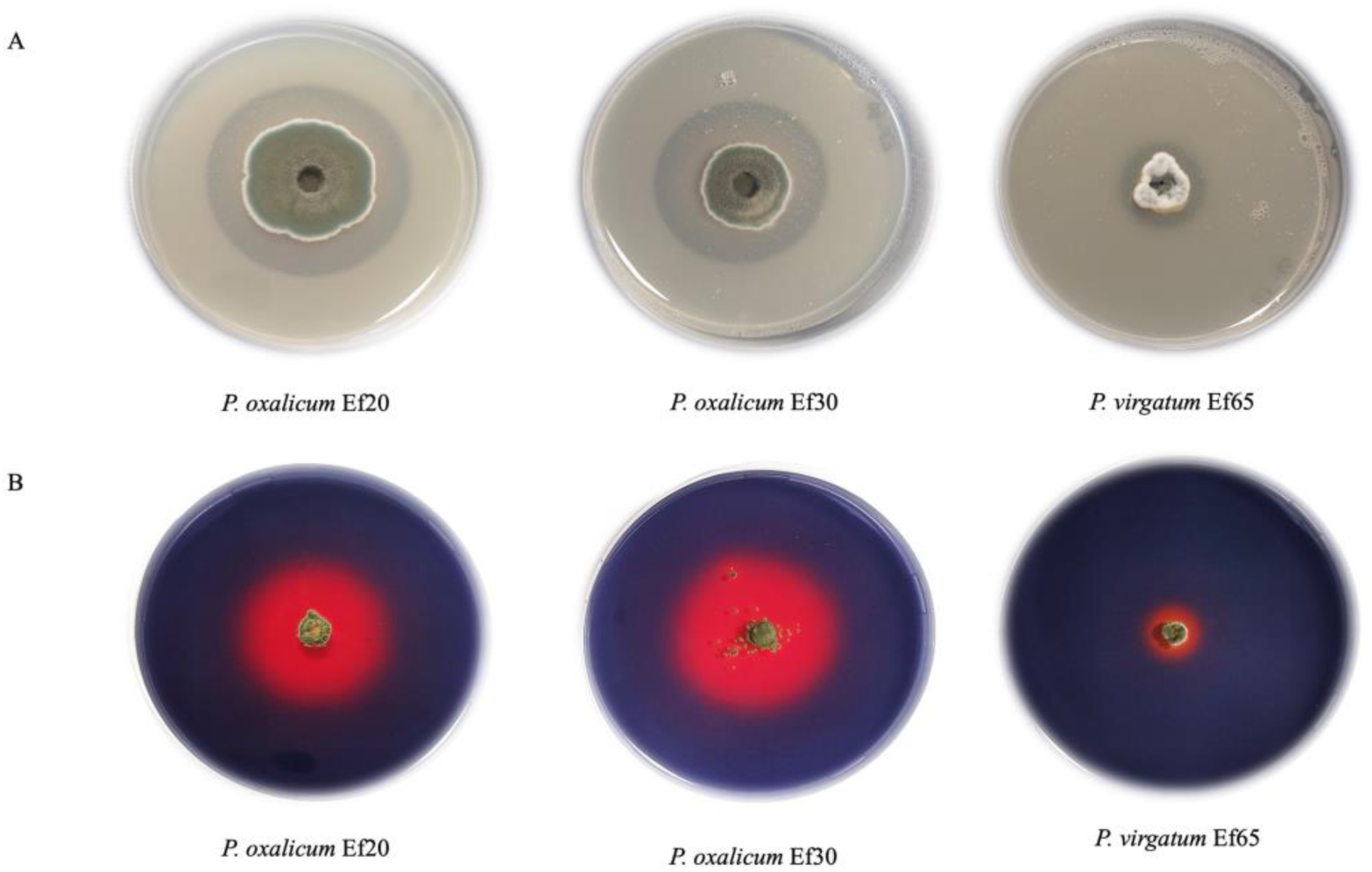
Plant growth-promoting traits: Phosphate solubilization and siderophore production of different endophytic fungal isolates *P. oxalicum* Ef20, *P. oxalicum* Ef30 and *P. virgatum* Ef65. Pikovskaya agar is shown by a clear zone (halo) around the colony after 3 days of incubation (A). Siderophore production was observed in the colour change of the medium from blue to orange on CAS agar plates (B). The experiment was repeated three times with three replications.

**Table 1.** Recognition of plant growth-promoting traits of different endophytic fungal isolates. Extracellular enzymatic activity, NH_3_ production of different endophytic fungal isolates of *P. oxalicum* Ef20 and Ef30 and *P. virgatum* Ef65. ‘+++’, strong producer of ammonia and lytic enzymes ‘++’ moderate; ‘−’, indicates a negative result. Phospate solubilisation, Siderophore and IAA production, values are the mean ± SD (n=3). Different lowercase letters (a, b, and c) denote significant differences at *P* ≤ 0.05 analyzed by Tukey’s test in Graphpad Prism. Three independent experiments were performed each with three replicates.

| Endophytic fungal Isolates | Phosphate solubilization (clear zone diameter in mm) | Siderophore production (clear zone diameter in mm) | IAA ($\mu\text{g} / \text{mL}$) | $\text{NH}_3$ production | Lytic enzymes | | |
| --- | --- | --- | --- | --- | --- | --- | --- |
|  |  |  |  |  | Protease | Cellulase | Lipase |
| *Penicillium oxalicum* Ef20 | 28.33 ± 0.58<sup>b</sup> | 30.7 ± 1.15<sup>b</sup> | 103.94 ± 2.69<sup>a</sup> | +++ | +++ | +++ | ++ |
| *P. oxalicum* Ef30 | 34.67 ± 0.59<sup>a</sup> | 35 ± 2.64<sup>a</sup> | 104.38 ± 3.40<sup>a</sup> | +++ | +++ | +++ | ++ |
| *P. virgatum* Ef65 | 11.67 ± 0.62<sup>c</sup> | 17.33 ± 0.57<sup>c</sup> | 31.28 ± 1.01<sup>b</sup> | +++ | + | + | - |

### 3.3. *P. oxalicum* isolates inhibit and disrupt fungal phytopathogen growth *in vitro*

The antagonistic potential of the three selected endophytic fungal isolates (*P. oxalicum* Ef20 and Ef30, and *P. virgatum* Ef65) was tested against phytopathogenic fungi that can cause disease in barley: *R. collo-cygni*, *Pyrenophora teres* and *Fusarium graminearum* (Fig. 3). Ef30 had a strong inhibitory effect on *R. collo-cygni* growth, producing an inhibition zone of 27 ± 1.47 mm, which was significantly higher inhibition than with Ef20 (20± 2.70 mm) and Ef65 (3.00 ± 0.58 mm) (Fig. 3A, D). Furthermore, Ef30 exhibited strong radial mycelia inhibition of *P. teres* (79 ± 1.57 %) (Fig. 3B, E) and *F. graminearum* (57.14 ±1.65%) (Fig. 3C,F) which was significantly higher than the radial growth inhibition observed for Ef20 against *P. teres* (72 ±1.73%) and *F. graminearum* (52.10 ±0.89 %). Both Ef20 and Ef30 showed significantly greater inhibition of radial growth compared to Ef65 for *P. teres* (40 .0± 1.65%) and *F. graminearum* (39.5 ± 1.37%).

**Fig. 3.**
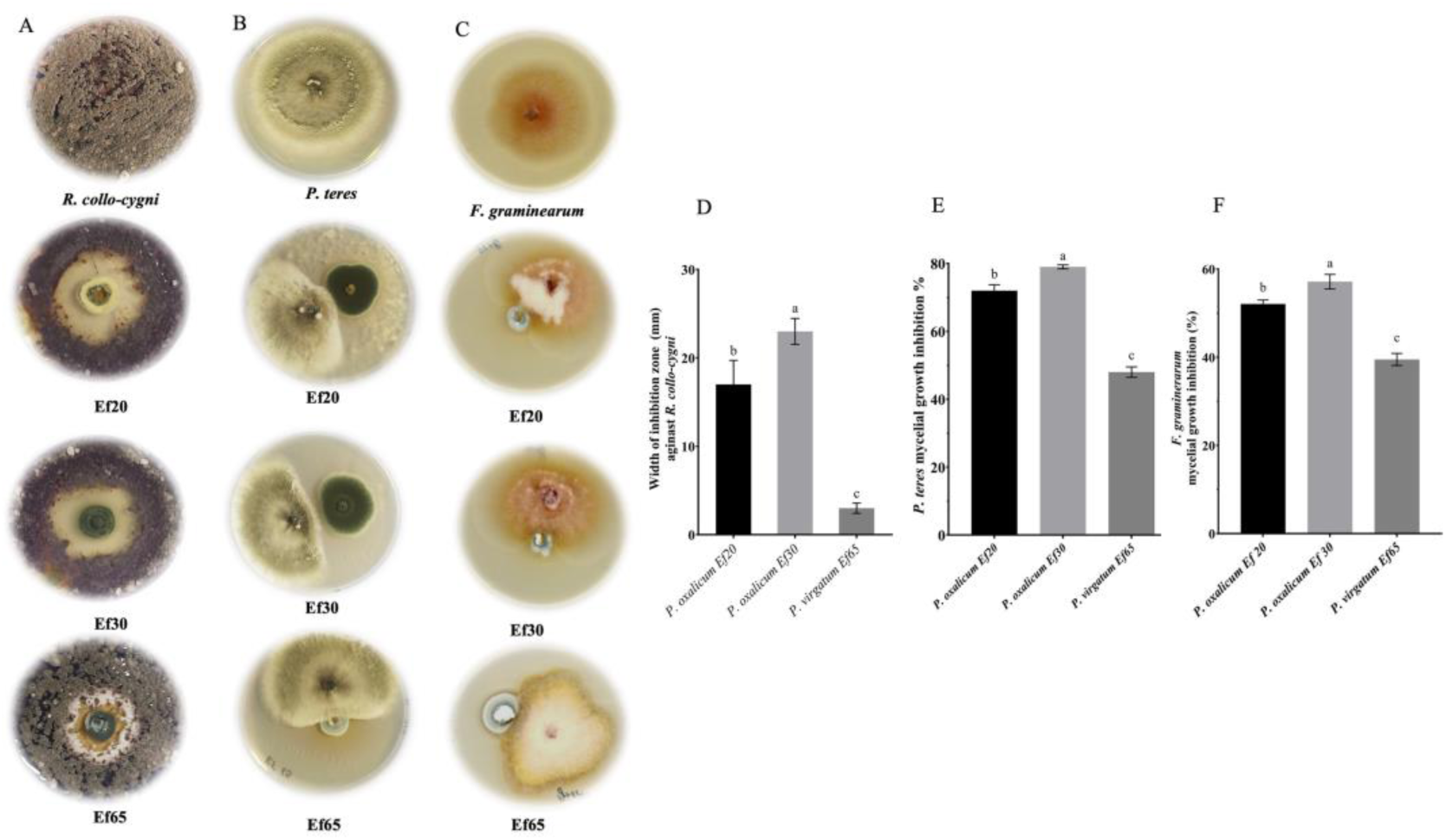
Confrontation assays on PDA between fungal endophytes *Penicillium oxalicum* Ef20, Ef30 and *P. virgatum* Ef65 against three fungal pathogens of barley. PDA plates with the fungal pathogens *(R. collo-cygni, P. teres* and *F. graminearum)* with and without the endophytic fungi are shown after 7 and 4 days. (A-C). For *R. collo-cygni, the* width of the zone of inhibition was measured in diameter (mm) (D). The percentage of *P. teres* and *F. graminearum* inhibition induced by each strain compared to the control (E,F); The data in the figure are mean ± SD; different letters represent statistically significant differences *P* ≤ 0.05). Three independent experiments were performed, each consisting of three replications.

To study inhibition of fungal pathogen growth by the endophytic fungi, samples were collected from the boundary zone of inhibition (the area around the fungi where pathogen hyphal growth is restricted). In the presence of the endophytic fungi (Ef20, Ef30 or Ef65), the hyphae of *R. collo-cygni*, *P. teres* and *F. graminearum* underwent several morphological changes including irregular, distorted, shrivelled and in some cases vacuolated hyphae (Fig. S3 D-L). On the contrary, the hyphae of *R. collo-cygni, P. teres,* and *F. graminearum* appeared smooth, uniform and undisrupted, with new hyphal buds arising on control plates without the endophytic fungi (Fig. S3 A-C).

### 3.4. *P. oxalicum* antifungal activity present in the culture filtrate, protects barley plants from *RLS*

To determine whether the growth inhibition of *R. collo-cygni* by *P. oxalicum* endophytes was due to secreted compounds in the media, endophytes were grown alone in liquid media for 7 days, after which the fungal biomass was removed leaving only the culture filtrate. *R. collo-cygni* growth was significantly inhibited by the culture filtrate of *Penicillium oxalicum* Ef20 and Ef30 (Fig. S4 B-C) compared to control plates (Fig. S4A), while no inhibitory effect was found with the *P. virgatum* culture filtrate Ef65 (Fig. S4D).

Based on the antifungal activity of the culture filtrate against *R. collo-cygni,* the potential to inhibit disease symptoms of *Ramularia Leaf Spot* was investigated *in planta*. Fourteen-day-old barley seedlings were inoculated with *R. collo-cygni* and, after 48 hrs, subsequently sprayed with the culture filtrate from *P. oxalicum* Ef30. The following conditions were tested: (i) *R. collo-cygni* inoculation only (ii) *R. collo-cygni* inoculation with the culture filtrate application, (iii) culture filtrate applied only, (iv) tween control (Tween 20). Barley seedlings that had not been sprayed with the culture filtrate had significantly higher RLS disease symptoms than seedlings sprayed with the culture filtrate (*P* ≤0.05) (Fig. 4A). On average, the culture filtrate significantly reduced RLS disease symptoms (Fig. 4). Barley that were not inoculated with *R. collo-cygni* did not exhibit disease symptoms. These results indicate that spraying with the Ef30 *P. oxalicum* culture filtrate protects barley from RLS disease (Fig. 4).

**Fig. 4.**
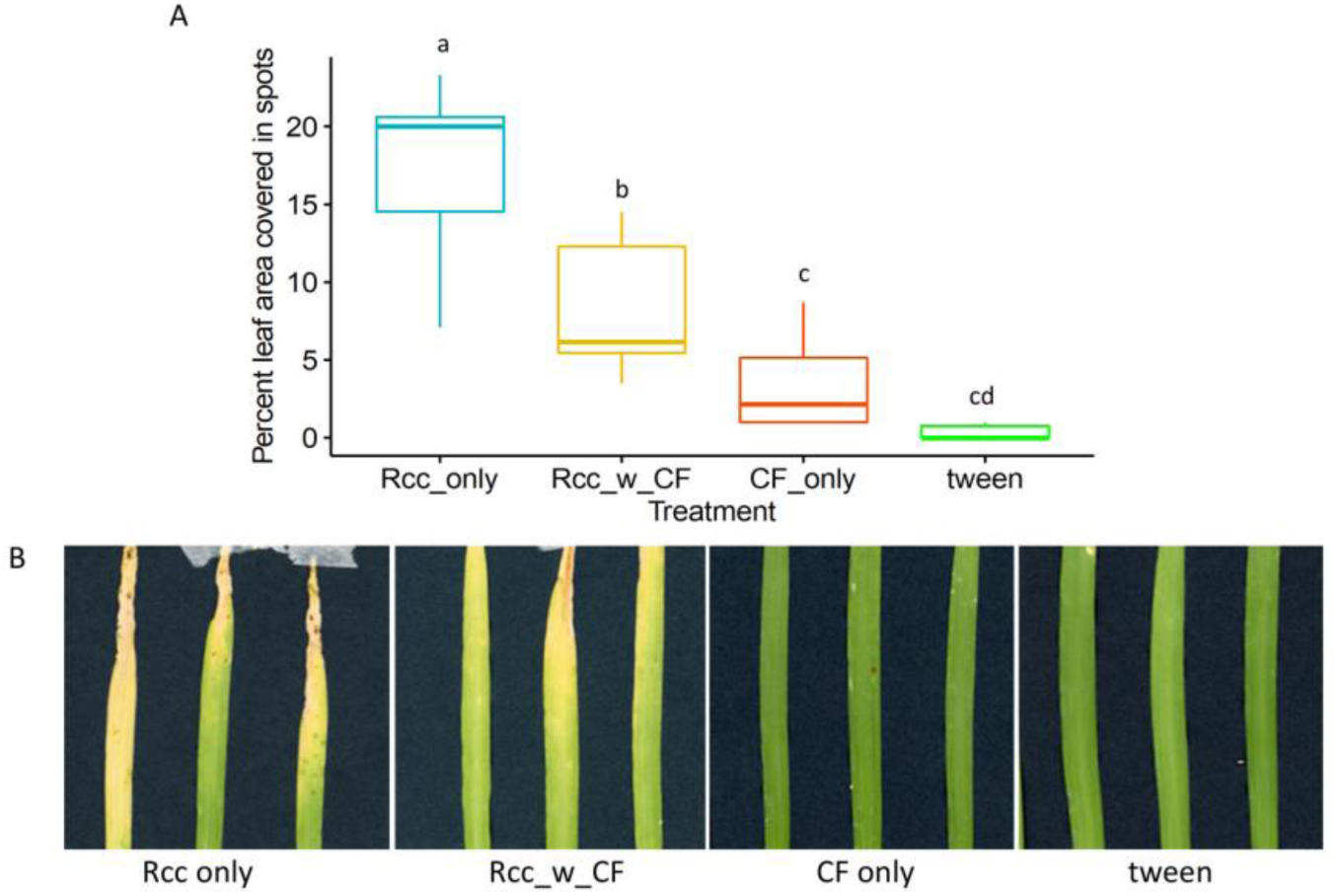
Percentage of leaf area covered in spots associated with RLS symptoms. (A). Barley leaves show symptoms for each treatment 17 dpi (B). Representative of three independent experiments. Different letters represent statistically significant differences (*P* ≤0.05).

### 3.5. Extraction and identification of an antifungal compound, Pyrrolidone-5-carboxylic acid from the *P. oxalicum* aqueous phase

The culture filtrate of *P. oxalicum* (Ef20, Ef30) *P. virgatum* (Ef65) and *P. verruculosus* (Ef39), which did not inhibit *R. collo-cygni* growth in the confrontation assay was collected as a negative control. Following extraction using an equal volume of ethyl acetate (1:1). The organic phase (upper ethyl acetate phase) and the aqueous phase (lower phase) were separated and subsequently lyophilized. The ethyl acetate phase was resuspended in methanol (based on solubility), and the aqueous phase was resuspended in sterile deionised water. Both the ethyl-acetate and aqueous extracts were tested for antifungal activity against *R. collo-cygni*. Only the aqueous phase applied on a filter disk exhibited antifungal activity against *R. collo-cygni* (Fig. S5), indicating that the active metabolites are present only in the aqueous phase. The ethyl acetate and aqueous extracts from the control endophytic fungus Ef39 did not exhibit inhibitory effects against *R. collo-cygni*.

A metabolomic approach using LC-MS/MS was performed to elucidate the composition of the active antifungal compound present in the aqueous extract of *P. oxalicum* Ef30 and Ef20 by comparison with *P. verruculosus* Ef39 and *P. virgatum* Ef65. The aqueous extract was analysed using untargeted metabolomics LC-MS/MS in both the negative and positive mode of ionization of the mass spectrometer. A list of metabolites was generated, and the compound mass was compared to the MELTIN database for each culture filtrate.

Based on the compound mass and the retention time, 18 metabolites were identified as common to Ef20 and Ef30, which were absent in Ef65 and Ef39 (Table S2, Fig. 5A ). Among the secondary metabolites identified from the literature as potential antifungals were 2-Pyrrolidone-5-carboxylic acid, 3-Methyl-2-oxovaleric acid, L,L-Cyclo (leucylprolyl), Mevalonolactone, and Pyridoxine (Table S3) [55,56,57,58]. Further analysis, by comparing the mass spectra with the MS spectral database (MELTIN database), confirmed the chemical compound as 2-Pyrollidone-5-carboxylic acid (2Py-5CA) based on the data of molecular formula, exact molecular mass, and retention time (Fig. 5B) match with the peaks of the 2-Pyrollidone-5-carboxylic acid standard (Fig. 5C). The compound 2-Pyrrolidone-5-Carboxylic (t_R_ = 1.14) displayed molecular ion [M^+^] protonated molecule [M+]^+^ at m/z 80.04 and 130.05, respectively.

**Fig. 5.**
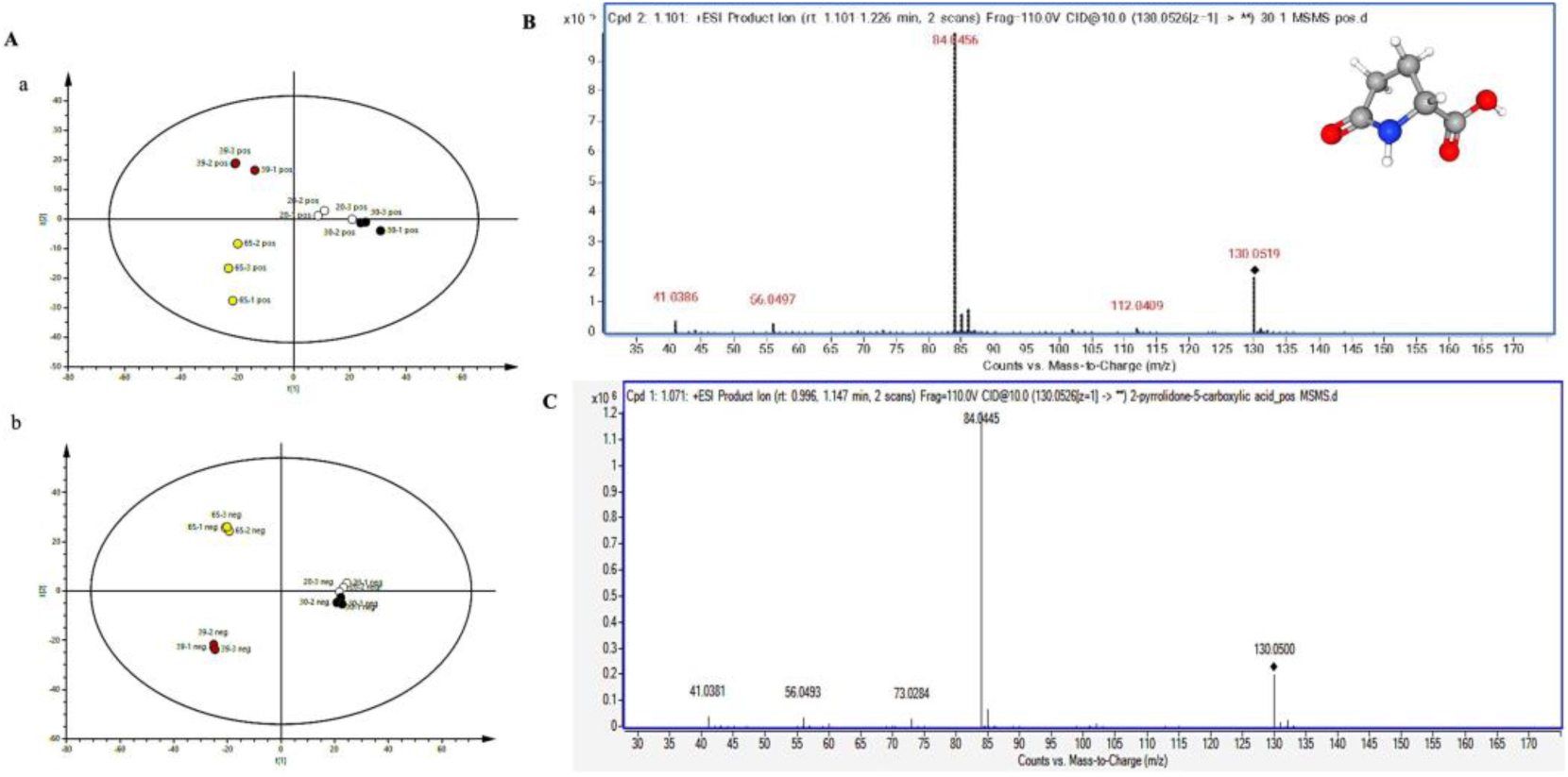
Principal component analysis of metabolites in aqueous fraction at positive (a) and negative (b) mode for Ef20, Ef30, Ef39, and Ef65. Pool of compounds based on sharing the same molecular formula in the culture filtrate of Ef20 and Ef30. Whereas no activity has been shown by the Ef39 and Ef65 culture filtrate (A). Mass spectra of 2-Pyrollidone-5-carboxylic acid and corresponding adducts detected in the endophytic fungal extract isolate *Penicillium oxalicum* Ef30 and Ef20. The detection was monitored at MS-ESI (+) spectroscopy at a probe temperature of 350 °C. (B) The fragments of 5-Pyrrolidone-2-Carboxylic acid (2Py-5CA) and (C) standard matched with MELTIN database at 130.0526 m/z in positive mode. 3D-structure retrieved from NCBI PubMed database 2-Pyrrolidone-5-carboxylic acid (B).

### 3.6. Pyrrolidone-5-carboxylic acid (2Py-5CA) inhibits the growth and development of fungal pathogens

*R. collo-cygni* and *Z. tritici* were evenly spread on PDA with different concentrations of 2Py-5CA (10, 20, 40, 80 and 100 µg/mL) to evaluate antifungal activity. PDA plates amended with 0.5% DMSO were used as a control. The percent surface area covered by *R. collo-cygni* was 43.20 ± 4.77 at 10 µg/mL 2Py-5CA , 36.61% ± 3.10 at 20 µg/mL 2Py-5CA, compared to 19.92% ± 3.42 at 80 µg/mL 2Py-5CA (Fig. 6A, C). Similarly, for *Z. tritici* the surface area covered was 61.46 ± 3.67 at 10 µg/mL 2Py-5CA, 43.48%±3.23 at 20 µg/mL and 26.64 %± 2.93 at 80 µg/mL 2Py-5CA (Fig. 6B, D). The results indicate a significant inhibition of growth against *R. collo-cygni* and *Z. tritici*, demonstrating a strong inhibitory effect by compound 2Py-5CA. This finding underscores the potential effectiveness of the treatment in controlling these pathogens.

**Fig. 6.**
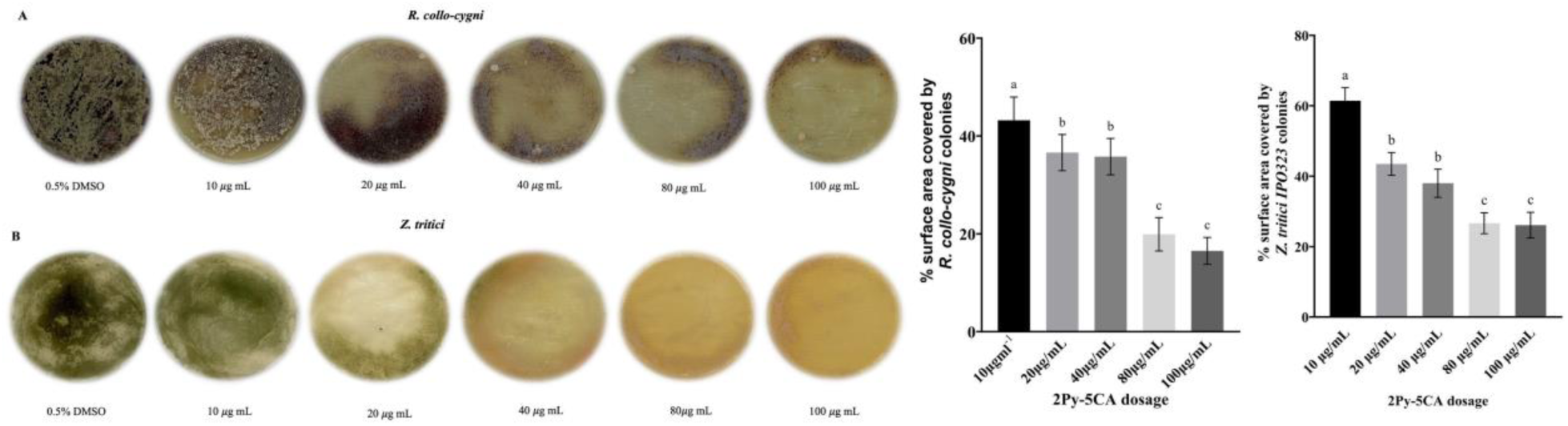
Antifungal effect of 5-Pyrrolidone-2-Carboxylic acid (2Py-5CA) on the growth of *R.collo cygni (RCC*) (A) and *Z. tritici*. (B) PDA with 10 *μ*g /mL^−1^-100 *μ*g/mL^−1^ of 2Py-5CA after 6 days of incubation, 0.5% DMSO served as a control. Percent (%) surface area covered by growth of *RCC* (C) and *Z. tritici* (D) was measured using ImageJ (https://imagej.net). The data in the figure are the mean ± SD; different letters represent statistically significant differences (*P* ≤ 0.05). Three independent experiments consisted of three replications with each treatment including controls (0.5% DMSO).

The impact of 2Py-5CA on the hyphal growth of the pathogens *F. graminearum, P. teres* and *B. cinerea* at different concentrations (10 - 100 µg/mL) was also examined with 0.5% DMSO as a control (Fig. 7). The results are shown as the percentage radial growth inhibition (PRGI) of the pathogen on PDA. A mycelial plug (5 mm), from the perimeter of a six-day-old actively growing colony of *P. teres*, *F. graminearum* and *B. cinerea* was placed in the centre of a PDA plate. All the petri dishes were incubated for 3 days containing fungal pathogens (*P. teres*, *F. graminearum* and *B. cinerea*) at 28 °C. The results show that 2Py-5CA inhibited the hyphal growth of these phytopathogens by reducing the radial hyphal diameter. At 20 µg/mL, the inhibition of radial growth for *F. graminearum* (Fg) was 31% ± 1.99%, compared to a maximum inhibition of growth of 56% ± 2.00 % at 80 µg/mL (Fig. 7A,D). Similarly, *Botrytis cinerea* on 20 µg/mL and 80 µg/mL 2Py-5CA caused 23.31% ± 1.13% and 46.65% ± 1.92 % mycelial radial growth inhibition respectively, compared to the 0.5% DMSO control (Fig. 7B,E). For the fungal pathogen *P. teres* the inhibition of hyphal growth was 22%±1.00% at 1 µg/mL 2Py-5CA, whereas, at 20 µg/mL, 76.3% ± 1.53% inhibition of the hyphal growth was found for 2Py-5CA (Fig. 7C,F). These results demonstrate a broad-spectrum biofungicidal nature of 2Py-5CA which is produced by *P. oxalicum* Ef30 against fungal pathogens.

**Fig 7.**
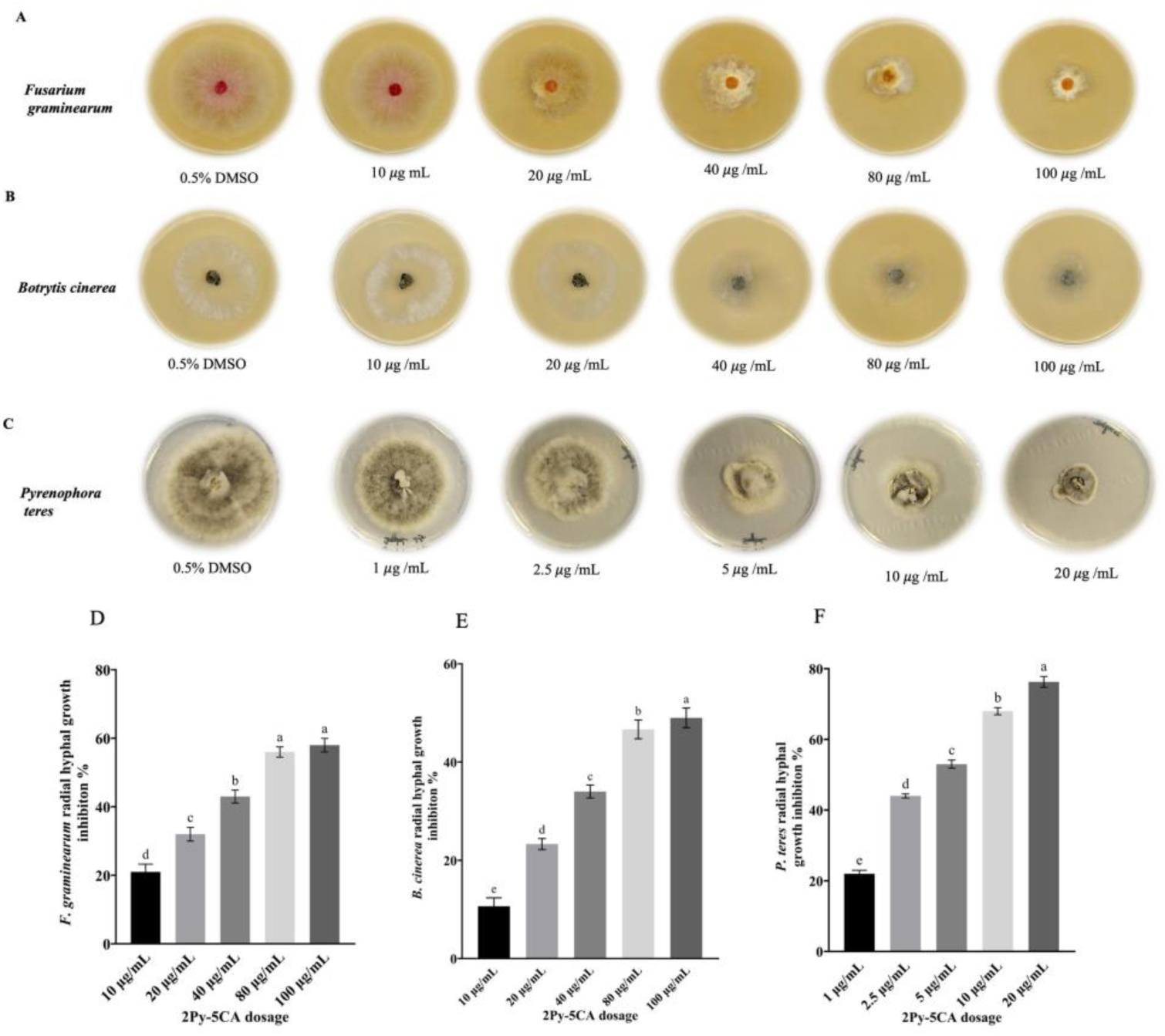
Antifungal effect of 2-Pyrrolidone-5-carboxylic acid (2Py-5CA) on the growth of *F. graminearum*, *B. cinerea* and *P. teres.* PDA with 0.5% DMSO served as a control (top left and reduction in growth treated with different concentrations (10*μ*g /mL-100 *μ*g/mL) of 2Py-5CA and for *F*. *gramineraum*, *B. cinerea* and *P.teres* (1*μ*g/mL). Inhibition of fungal pathogen growth by 2Py-5CA represented as the inhibition percentage (%) in the diameter of the radial growth of the fungal mycelia in the treated plates compared to that of the control PDA plates; *F. graminearum* (A, D), *B. cinerea* (B,E) and *P. teres* (C,F). The data in the figure are the mean ± SD; different letters represent statistically significant differences one way ANOVA Tukey’s tests (*P* ≤ 0.05). The experiment was repeated three times independently with three replicates.

Light microscopy was used to visualize the impact of 2Py-5CA on the morphology of *R. collo-cygni, P. teres, Z. tritici*, and *F. graminearum* (Fig. 8). Treatment with 2Py-5CA was responsible for aberrant morphological changes and anomalies of the pathogenic fungal hyphae and spores. The hyphae in the untreated control group have a regular smooth surface (Fig. 8A-C) and a high density of *Z. tritici* blastospores (Fig. 8D). In contrast, the fungal pathogen mycelia treated with 2Py-5CA are associated with deformed, expanded and disintegrated hyphae. The ends of the mycelia were swollen, with round hyphal tips and conglobate structures (Fig. 8E-H). Furthermore, deformation of *F. graminearum* macroconidia morphology, bending, tortuous growth and evident swelling following treatment with different concentrations of 2Py-5CA were observed (Fig. 9 B-E) when compared with control plates (0.5% DMSO), which displayed smooth, uniform structures with well-defined septa (Fig. 9 A). Thus, 2Py-5CA changes the morphology of hyphae and spores inhibiting the growth of the fungal pathogens tested *in vitro*.

**Fig. 8.**
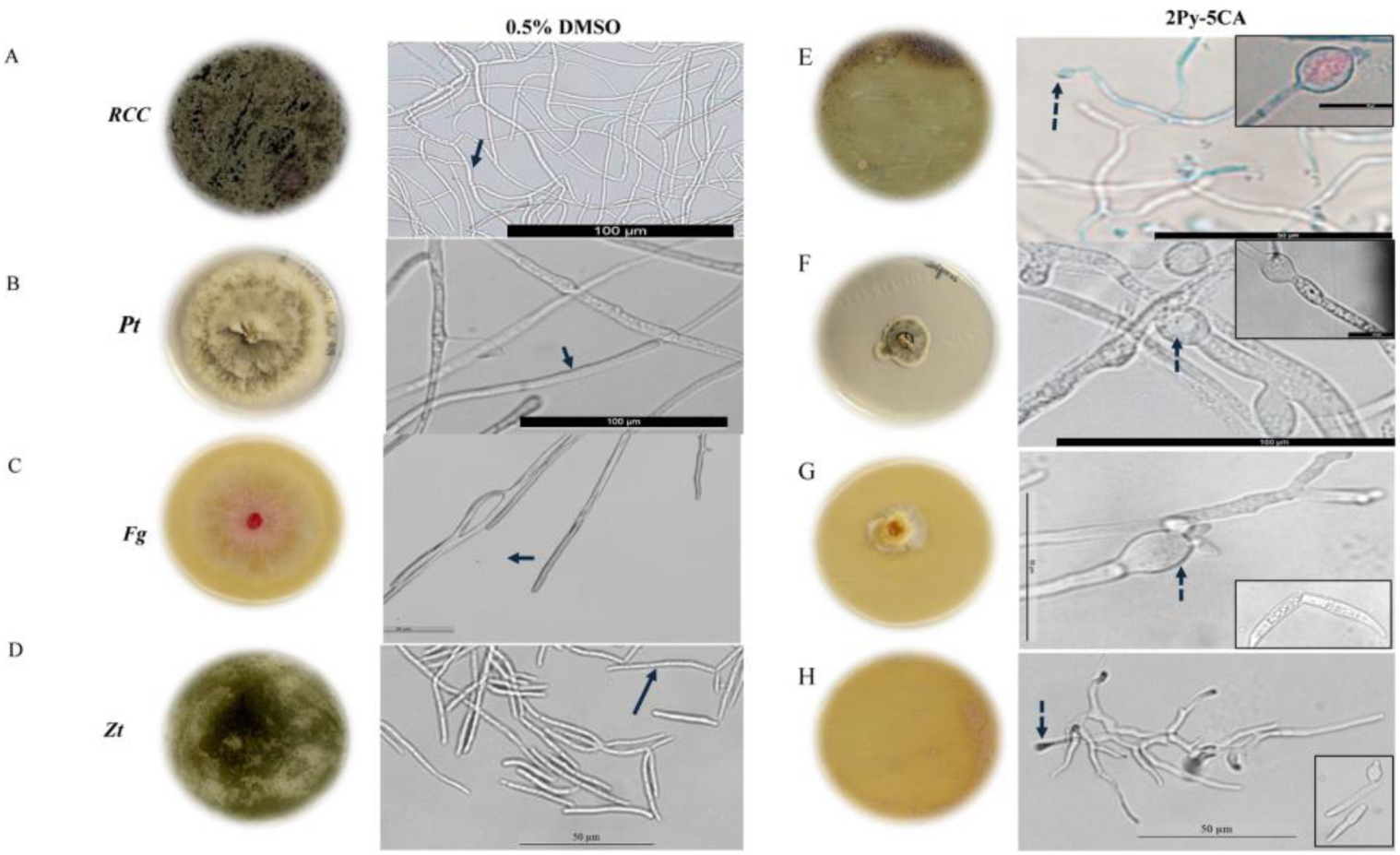
Effects of 2Py-5CA on hyphae of *R. collo-cygni, Z. tritici*, *F graminearum* and *P. teres*. In the controls (0.5% DMSO) pathogens (*RCC, Pt* and *Fg*) hyphae are undisrupted, smooth (A, B, C) and have a high density of blastospores of *Z. tritici* (D) (full black arrow); whereas plates with compound 2Py-5CA dosage at (80 µg/ml^−1^) show hyphal deformation swelling, ball and round projections of hyphal tip (E, G, H), conglobate structure (E-F), was observed (dashed arrows head). Inset images showed bulging type morphology of hyphae (E, F, H) and broken macroconidia (inset image) (G), suggesting a fungicidal mode of action. Scale bar = 100 and 50 *μ*m.

**Fig. 9.**
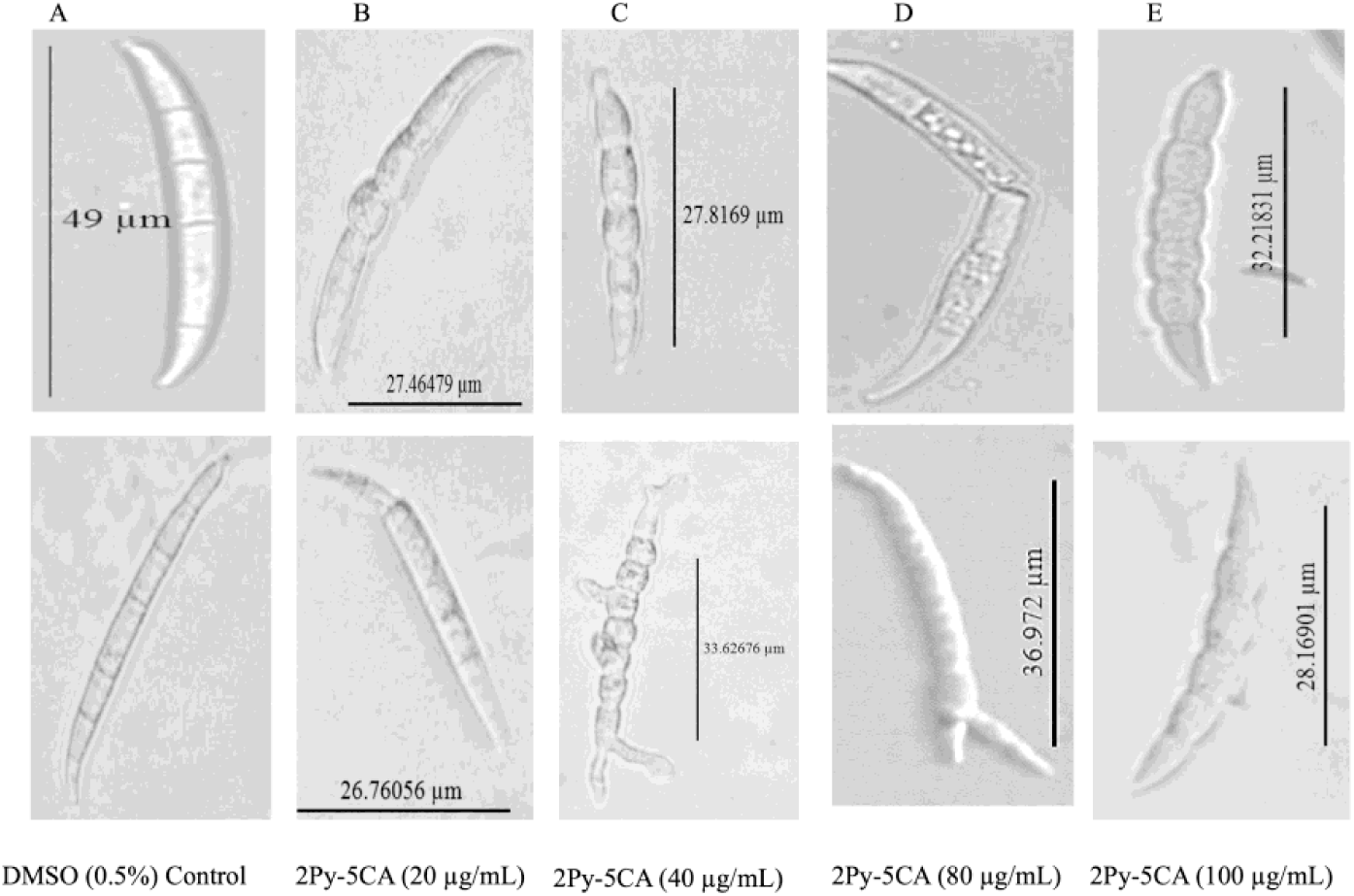
Toxicity of 2Py-5CA dosage on *Fusarium graminearum (Fg)* spores. Macroconidia of *Fg* on 0.5% DMSO controls are elongated, smooth, uniform with a remarkable septa (A). Distorted morphology with tortuous growth, evident swelling, broken macroconidia and different size was observed when treated with different 2Py-5CA dosages (B-E) (20 µg/mL - 100 µg/mL ).

### 3.7. Control of RLS in barley by 2-Pyrroldione-5-carboxylic acid (2Py-5CA)

The control efficacy of 2Py-5CA against RLS was evaluated *in planta* (Fig. S6). Barley seedlings were first inoculated with a *R. collo-cygni* suspension followed by spraying with different concentrations of 2Py-5CA (10 µg/mL, 20 µg/mL, 40 µg/mL, and 80 µg/mL) at 48 hours post inoculation (hpi) as depicted (Fig. S6). Control barley plants were sprayed with 0.5% DMSO. Disease scoring was performed on the second leaf, recording the relative area of the leaf covered in lesions. The control plants sprayed with 0.5% DMSO had significantly higher RLS disease symptoms (4.98 ± 1.2)compared to plants sprayed with 80 µg/mL 2Py-5CA (3.60 ± 0.77) (*P* ≤ 0.04) (Fig. 10). In contrast, the plants sprayed with the moderate concentration of compound 2Py-5CA at 20 µg/mL concentration leaves also had a significant reduction of RLS symptoms (2.53 ± 0.52) compared to 0.5% DMSO alone (*P* ≤ 0.001) and 80 µg/mL 2Py-5CA (*P* ≤ 0.02) (Fig.10). This *in planta* assays suggest that a moderate concentration (20 µg/mL) of 2Py-5CA is effective at reducing RLS disease symptoms in barley.

**Fig. 10.**
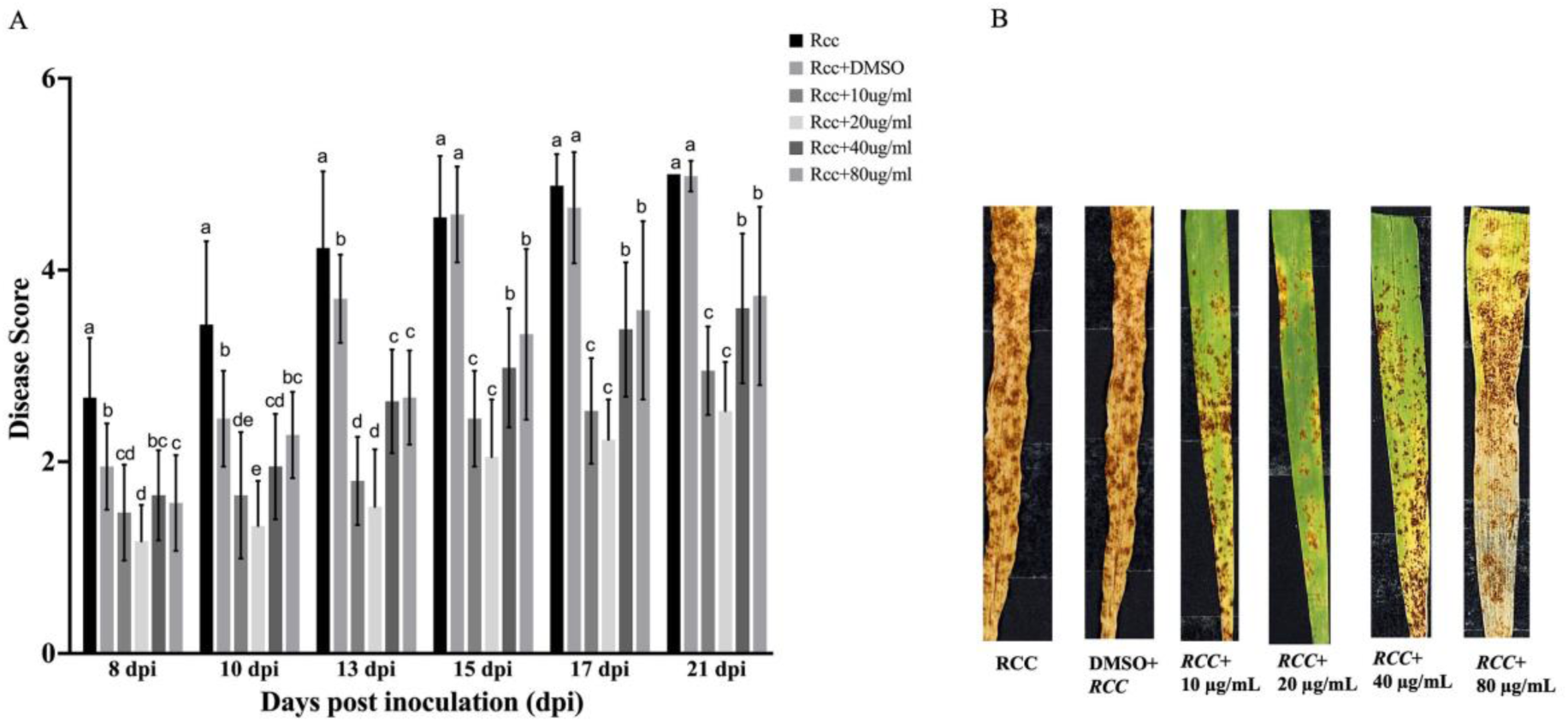
Reduction of Ramularia leaf spot (RLS) symptoms on barley leaves treated with 2Py-5CA. Disease severity of barley plants pre-treated with pathogen *RCC* and 48 hrs, thereafter, treated with different doses (10 - 80 µg/mL) of 2Py-5CA. *RCC* symptoms were assessed from 8 dpi up to 21 dpi. DMSO 0.5% served as a control (A). Different letters (a-f) indicate significant differences among different treatments one-way ANOVA followed by Tukey’s tests *P* ≤ 0.05 indicates significant differences between treatments. Images of RLS disease reduction after post-treatment with 2Py-5CA dosage at 10 – 80 µg/mL. Pretreatment with *RCC* 48 hours before 2Py-5CA and control inoculation with DMSO (0.5%) used as a negative control (B). Images show the RLS symptoms after 21 days post-inoculation (21 dpi). Three independent experiments were performed with two replications (10 leaves each from individual seedlings per treatment per replicate).

### 3.8. Prediction of the secondary biosynthetic gene clusters (BGCs) which produce 5Py-2CA

The assembled genome sizes were 7,299,882bp for Ef30; 7,433,734 for Ef20, and 8,994,090 bp for Ef39. The final assemblies consisted of 610 contigs (Ef20), 893 contigs (Ef30), and 714 contigs (Ef39), with N50 values of 557,391 bp, 189,033 bp, and 1,585,510 bp, respectively. The average GC content was 50.56% in Ef20, 50.5% in Ef30, and 46.4% in Ef39, reflecting moderate variation across the fungal genomes. AntiSMASH was used to predict secondary metabolite biosynthetic gene clusters (BGCs) in *P. oxalicum* strains Ef20, Ef30, and *P. verruculosus* Ef39, serving as a comparison. The predicted secondary metabolite (SM) gene clusters of strains Ef20 and Ef30 are characterised by ‘backbone enzymes’ that form the carbon skeleton of putative SMs including; non-ribosomal peptide synthase (NRPS), polyketide synthase (PKS), and terpene synthase. A total of 52, 41 and 63 secondary metabolite BGCs were identified in Ef20, Ef30 and Ef39, respectively. These inlcude NRPS/NRPS-like enzymes, PKS/T1-PKS/ T3-PKS/ PKS-like enzymes, and terpene/terpene precursors, along with an NRPS-Indole synthetase, Fungal RiPP-NRPS, NI-siderophore and Isocyanide/ isocyanide-NRP gene clusters (Table S4). Six BGCs share gene homologies with known clusters in the MIBiG database present for endophytic fungal isolates Ef20 and Ef30, which were not present in Ef39 (Table S4). Therefore, Ef20 and Ef30 share common BGCs for bioactive compounds such as Atpenin B, Oxaleimide C, Aurofusarin, Histidyltryptophanyldiketopiperazine, Chaetoglobosin A and Paraherquamide (Table S5).

Further, AntiSMASH analysis results revealed a hybrid NRPS-indole gene cluster located in the regions 19.1 and 30.1 of strains Ef20 and Ef30 which exhibits 100% BLAST sequence similarity to the Paraherquamide (Phq) BGC from *Penicillium fellutanum* (GenBank: JQ708195.1) (Fig. 11A, B). Annotation *via* minimum information biosynthetic gene cluster (MIBiG) procluster confirmed that all fifteen genes within these regions are homologous to our query sequence. Those involved in the different analogs of the paraherquamide pathway in the MIBiG database (Fig. 11 C,D) and the paraherquamide structure contains a 2-pyrroldione core ring (Fig. 11E). Paraherquamide biosynthesis is initiated at the 2-Pyrrolidone moiety, leading to structural divergences among paraherquamide analogs through various enzymatic modifications. The paraherquamide biosynthetic gene cluster (BGC) includes a gene encoding L-1-pyrroline-5-carboxylate reductase, an enzyme pivotal in the L-proline biosynthetic pathway. This enzyme facilitates the conversion of 2-pyrrolidone-*di*-carboxylic acid (2Py-*di*-CA), a proline intermediate in the L-proline KEGG pathway (Fig. 12).

**Fig. 11.**
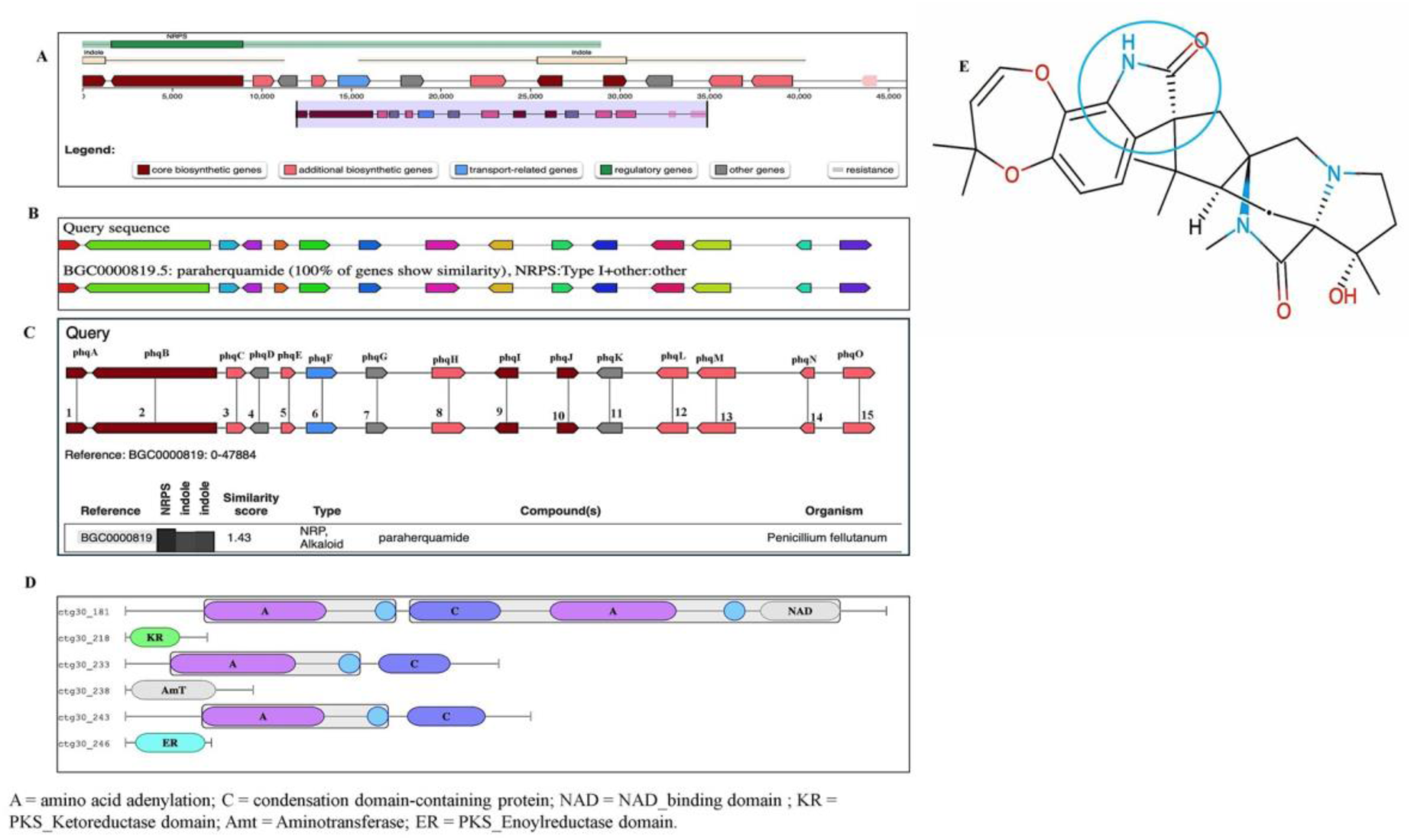
Hybrid NRPS-Indole biosynthetic gene cluster in *Penicillium oxalicum* EF20 and Ef30 (A). Common biosynthetic gene cluster (paraherquamide) with high similarity of 100% in *Penicillium oxalicum* Ef20 and Ef30 (**B**). The NRPS-Indole hybrid gene cluster in the genome of endophytic strains Ef20 and Ef30 shows a similarity score of 1.43 with the paraherquamide cluster in *Penicillium fellutanum* (GenBank: JQ708195.1), according to the MIBiG database. The hybrid NRPS-indole gene cluster in region 19.1 and region 30.1 of strain Ef20 and Ef30 genome encoded genes: 1,phqA, 9 phqI and 10, phqJ encoded genes (prenyltransferase); 2, phqB (NRPS); 3, phqC (2OG-Fe(II)-oxygenase); 4, phqD (pyrroldine-5-carboxylate reductase dimer); 5, phqE (short-chain dehydrogenase); 6, phqF (efflux pump); 7, phqG (negative regulator);8, phqH (oxidoreductase); 11, phqK (FAD monooxygenase); 12, 13, and 15 phqL, phqM and phqO (P450 monooxygenase). 14, phqJ (methyltransferase) (C). Represents the NRPS domains involved in NRPS biosynthesis based on Pfam annotation. A = amino acid adenylation; C = condensation domain-containing protein; NAD = NAD_binding domain ; KR =PKS_Ketoreductase domain; Amt = Aminotransferase; ER = PKS_Enoylreductase domain (D). MIBiG database download chemical structure of paraherquamide with the 2-pyrrolidone core ring (highlighted with a circle) (E ).

**Fig. 12.**
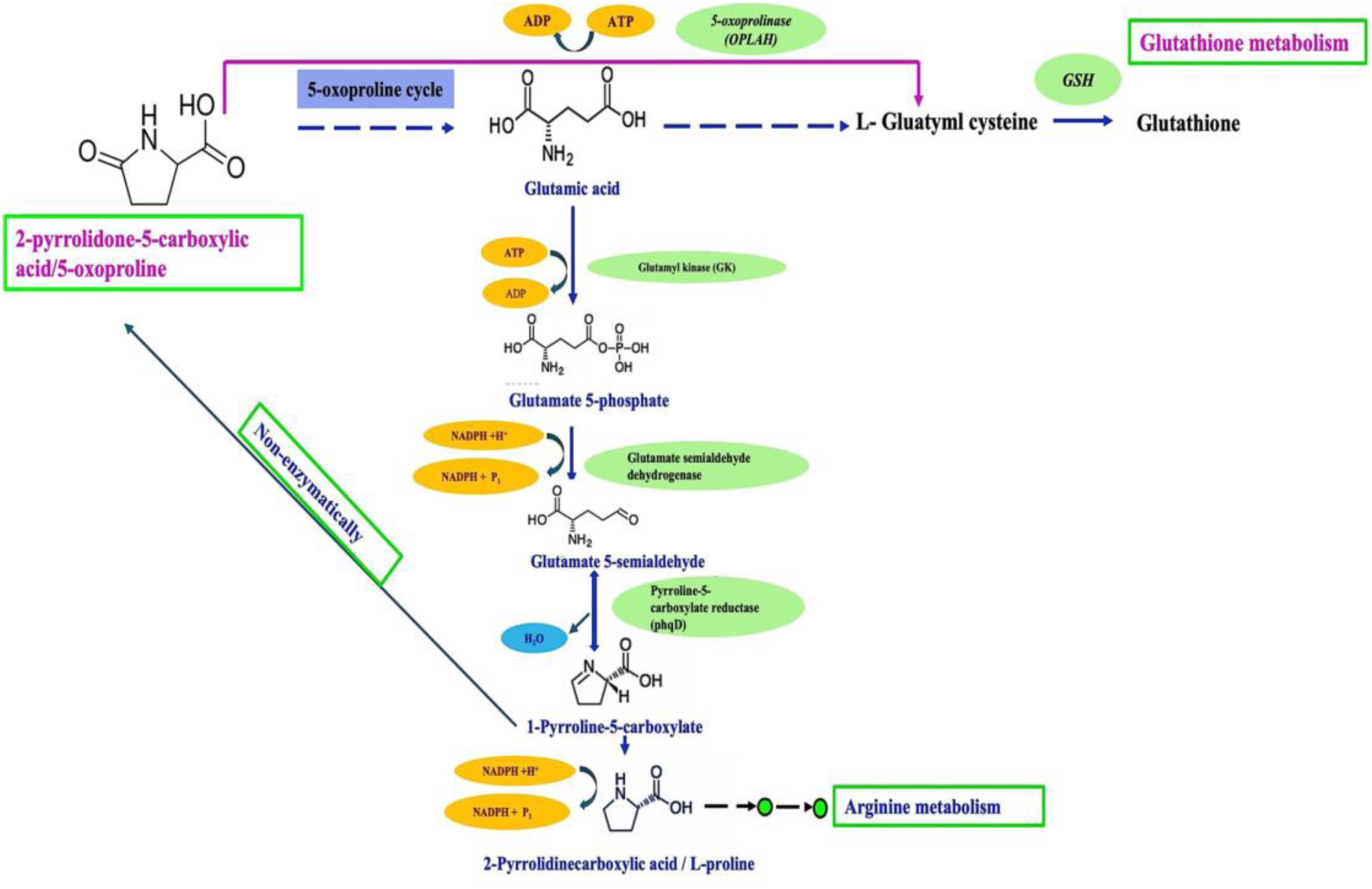
The biosynthetic pathway of L-proline and 2-pyrrolidone-di-carboxylic acid involves 1-pyrroline-5-carboxlate (P5C) as a crucial intermediate. L-Proline is derived from glutamate 5-phosphate *via* phosphorylation and dehydrogenation steps, leading to spontaneous cyclization into P5C which is then reduced to L-proline by an enzyme pyrroline-5-carboxylate reductase (5PCR) encoded by the phqD gene. A subsequent non-enzymatic reaction of 1-pyrroline-5-carboxlate between the carbonyl group of the carboxylic acid and the amino group of the pyrroline ring yields 5-oxoproline or, synonymously, 2-pyrroldione-5-carboxylic acid (2Py-5CA) as shown in the arrow. This 2Py-5CA is the structural derivative of L-proline and also a metabolite in the glutathione cycle.

The biosynthesis of proline begins with the methylation of isoleucine and subsequent reduction to methyl-pyrroline-5-carboxylic acid, which is then converted to L-proline (2-pyrrolidone-*di*-carboxylic acid) by the enzyme L-1-pyrroline-5-carboxylate reductase dimerization (P5CR dimer) encoded by *phqD* (Fig. 12). *PhqD* is part of a cluster essential for paraherquamide biosynthesis, a fungal indole alkaloid characterized by a 2-pyrrolidone core structure [59]. An enzymatic step involving glutamyl phosphate reductase (GPR) catalyzes the second reaction of the proline pathway, causing a reversal of its second biosynthetic pathway direction leading to the formation of 2-pyrrolidone-5-carboxylic acid (also known as 5-pyrrolidione-2-carboxylic acid), a structural analog of L-proline (Fig. 12) [60]. Further, KEGG analysis identified the involvement of *phqD*, which encodes for the 1-pyrroline-5-carboxylate reductase, in the arginine-proline-glutamate pathway supporting the biosynthesis of L-proline and its derivatives such as 2Py-*di*-CA (2-pyrrolidone-*di*-carboxylic acid) (Fig. S7).

## 4. Discussion

A major global challenge for crop production is yield losses of 10-23% due to fungal disease.^61^ The excessive use of chemical fungicides to prevent such losses has led to fungicide resistance. For example, there has been a rise in the resistance to major single-site fungicides, such as azoles and succinate dehydrogenase inhibitors (SDHIs) [[61], [62]] Prolonged intensive use of fertilisers and fungicides can lead to soil and aquatic life contamination, while exposure to fungicides during field spraying can pose health risks for farm workers [63–64]. Endophytic fungi, which reside within plant tissues, produce enzymes, secondary metabolites with antifungal, and plant growth-promoting (PGP) properties [65]. Biological control agents have gained considerable interest due to their environmental sustainability, cost-effectiveness, and diverse mechanisms of action.^66^ Several fungal genera, including *Penicillium spp.*, *Trichoderma spp.*, *Aspergillus spp.* and *Nigrospora spp.* have been identified as promising biocontrol agents [[67], [68], [69]].

Endophytic fungi support their host plants by enhancing nutrient availability under adverse climatic conditions, primarily through mechanisms such as phosphate solubilisation, siderophore production, and indole-3-acetic acid (IAA) synthesis [[70], [71], [72] ,[73]]. Endophytic fungi indirectly assist plants by generating compounds including ammonia and hydrolytic enzymes as well as engaging in antibiosis [74]. Our findings demonstrated that both endophytic strains of *Penicillium oxalicum* (Ef20 and Ef30) exhibit plant growth-promoting attributes including phosphate solubilation, siderophore, IAA, ammonia and lytic enzyme production. IAA is important for plant development and is considered a major phytohormone responsible for cell elongation, division, enlargement, and lateral root formation [[75], [76]]. Endophytic fungi also produce siderophores that enhance iron absorption and protect plants against infection. Previously, phosphate solubilisation was shown by the endophytic fungal isolates *Penicillium crustosum* EP-2, *P. chrysogenum* EP-3, and *Aspergillus flavus* EP-145 from the medicinal plant species *Ephedra pachyclada* [77]. These fungal endophytic isolates from *E. pachyclada* improved the growth of Maize (*Zea mays* L). *Penicillium citrinum* isolated from the leaves of wheat (*Triticum aestivum*) variety PBW 725 was able to produce IAA and ammonia [78]. While, *Penicillium radicicola* and *P. archidendri* have been reported for potential PGP activities for cultivating seedlings of Himalayan silver birch (*Betula utilis* D. Don) [79]. This study establishes that *P. oxalicum* (Ef20 and Ef30) have strong extracellular enzyme production, such as cellulase, protease, gelatinase and lipase. The hydrolytic enzymes cellulase, amylase, protease, and lipase, etc., have been widely used in crops (rice, soybean, wheat, and maize) for protection against invading plant pathogens [[80], [81], [82], [83]]. *Penicillium sp.* L3 isolated from the leaf of acai palms (*Euterpe precatoria*), also produces hydrolase enzymes that act against pathogens [81].

In our study, we identified two strains *P. oxalicum* Ef20 and Ef30 that have strong antifungal potential against phytopathogens of barley, including *R. collo-cygni*, *Pyrenophora teres* and *Fusarium graminearum in vitro*. Endophytic fungi *Penicillium cintrinum* DBR-9, isolated from the medicinal plant *Stehania kwangsiensis,* has antifungal activity against the oomycete *P. parasitica* var. *nicotianae*, fungi *Alternaria oleracea,* and *Bipolaris maydis* [84]. It was shown that polyketides produced by *Penicillium cintrinum* DBR-9 inhibit hyphal growth of *P. parasitica* var *nicotianae* [84]. Furthermore, *P. striatisporum* Pst10 has antagonistic effects against the oomycete *Phytophthora capsica,* which causes root rot in chilli pepper [85]. Polyketides isolated from *P. striatisporum* Pst10 induce morphological aberrations such as swelling and a noticeable loss of hyphal pigmentation [85]. *Penicillium oxalicum* was previously found to be antagonistic against the fungal pathogens *Fusarium oxysporum f. sp* and *Verticillium dahlia,* which cause disease in tomato and pea, respectively [[86], [87]]. *Penicillium oxalicum* also has an antagonistic effect against *Alternaria alternata* and was found to penetrate *A. alternata* hyphae to disintegrate the conidiophores [88]. The *Penicillium* genus is known for being a versatile producer of various secondary metabolites with antifungal, antibacterial, and antitumor toxins [[89], [90]]. Various species of *Penicillium* synthesise secondary metabolites such as Patulin, Ochratoxin A, Penitrem A, and Ochratoxin A (e.g *Penicillium rubens, P. flavigenum, P. dipodomyis, P. chrysogenum, P. nalgiovense, P. expansum, P. allii-sativi, P. verrucosum, P. commune, P. chrysogenum*) which exhibit larvicidal, inflammatory, antibacterial and antifungal activity [[91], [92], [93], [94]]. Altenusin and dehydroaltenusin, produced by two *Penicillium* species, show broad antifungal activity, inhibiting mycelial growth of several pathogens including *Fusarium solani*, *Fusarium oxysporum*, *Macrophomina phaseolina*, and *Cladosporium cladosporioides*. Altenusin and dehydroaltenusin, produced by two *Penicillium* species, show broad antifungal activity, inhibiting mycelial growth of several pathogens including *Fusarium solani*, *Fusarium oxysporum*, *Macrophomina phaseolina*, and *Cladosporium cladosporioides* [95].

We identified an acidic antifungal compound (2-pyrrolidone-5-carboxylic acid, 2Py-5CA), isolated for the first time in fungi, from the endophyte *P. oxalicum* that has fungicidal activity against cereal crop pathogens. Further, 2Py-5CA exhibits a broad spectrum antifungal activity against a range of cereal pathogens, *R. collo-cygni*, *F. graminearum*, *Z.tritici* and the fruit pathogen *B. cinerea in vitro*. Microscopic observations revealed that 2Py-5CA induced pronounced morphological abnormalities in fungal hyphae compared to untreated controls. The compound altered hyphal surface structure and tip integrity, resulting in distorted and irregular hyphal growth. These morphological deformations suggest that this may be part of the antifungal activity of 2Py-5CA. Previously, 2Py-5CA was found in the bacterial fermentation broth of the bacteria *Burkholderia* sp. HD05, isolated from Baimang River sediment, where 2Py-5CA was shown to inhibit the pathogenic oomycete *Saprolegnia sp*., which is important in aquaculture [96]. 2Py-5CA was found in bacterial strains of *Lactobacillus* and *Pediococcus* demonstrating antimicrobial properties effectively inhibiting common food spoilage bacteria such as *Enterobacter cloacae* 1575, *Pseudomonas fluorescens* KJLG, and *P. putida* 1560-2 [97]. Our *in planta* validation assay confirmed that treating barley seedlings with a moderate concentration (20 µg/mL) of 2Py-5CA reduced the progression of the economically important barley disease *RLS*. Results indicate that 2Py-5CA has a strong potential as a biocontrol agent and could decrease the need for chemical fungicides and help counter fungicide resistance, promoting sustainable agriculture practices. Previoulsy, the acidic antifungal metabolite Phenazine-carboxylic acid (PCA), derived from the culture filtrate of *Pseudomonas fluorescence* strain LBUM636, has been reported to suppress potato late blight caused by *Phytophthora infestans* [98]. *Penicillium cf. manginii* DSM 10449 produces the mycotoxin tetramic acid, and citreoviridin which inhibits the fungal pathogen *Hymenoscyphus fraxineus* [99]. Phenazine-1-carboxylic acid (PCA) acid produced by *Burkholderia* sp. HQB-1 bacteria exhibit antifungal activity against *Fusarium oxysporum* f. sp., *Colletotrichum gloeosporioides*, *Curvularia fallax*, and *Botrytis cinerea*. It also effectively protected banana plants from *Fusarium oxysporum* preventing *Fusarium* wilt [100]. A new phthalide derivative (-)-5-carboxylmellein and (-)-5-methylmellein were isolated from the endophytic fungus *Xylaria* sp. A novel series of indole-3-carboxylic acid derivatives has been reported to possess potent antibacterial activity against *Enterococcus faecalis* [101]. Also, short-chain derivatives of carboxylic acid have been shown to possess strong antifungal activity against *Fusarium oxysporum*, resulting in nearly 80% disease suppression in wheat seedlings using both *in vitro* and *in planta* experiments [102]. Also, it was found that the cinnamic acid-based carboxylated metabolites significantly disrupted the fungal cell wall of *Botrytis cinerea* under *in vitro* conditions [103]. These pyrrolidone derivatives exhibit diverse biological activities such as antiviral, antifungal, antibacterial, and anti-inflammatory properties [[104],[105], [106]].

AntiSMASH analysis provides insight into biosynthetic gene clusters and secondary metabolites relevant to discovering new antifungal metabolites present in the genome. *P. oxalicum* Ef20 and Ef30 contain various biosynthetic gene clusters (BGCs), such as PKSs, NRPSs, Hybrid NRPS-indole and Terpenes BGCs with potential for new secondary metabolites. NRPSs are crucial for producing peptides with antifungal and antibacterial activities [107]. We identified common biosynthetic gene clusters for Atpenin B, Oxaleimide C, Aurofusarin, Histidyltryptophanyldiketopiperazine, Chaetoglobosin A and Paraherquamide present only in *P. oxalicum* ( Ef20 and Ef30) but not in *Penicillium verruculosus* (Ef39). The paraherquamide biosynthetic gene cluster (BGC) includes a gene encoding L-1-pyrroline-5-carboxylate reductase, an enzyme pivotal in the L-proline biosynthetic pathway which facilitates conversion of 2-pyrrolidone-*di*-carboxylic acid (2Py-*di*-CA), a proline intermediate in the L-proline pathway. This enzymatic step not only links secondary metabolism with amino acid biosynthesis but also underscores the metabolic versatility encoded within the paraherquamide BGCs [[108], [109]].

## 5. Conclusion

This study reports, for the first time, the identification of an antifungal metabolite 2-Pyrrolidone-5-carboxylic acid (2Py-5CA), extracted from endophytic strains *P. oxalicum* Ef20 and Ef30 and exhibits broad-spectrum antifungal activityagainst cereal pathogens under *in vitro* conditions. Further, *in planta* assays have confirmed the biocontrol potential of 2Py-5CA, showing effective suppression of *Ramularia collo-cygni,* the causal agent of Ramularia leaf spot (RLS) in barley. Microscopic observation revealed hyphal disruption and condial deformation following treatment with a single purified compound, suggesting a direct impact on fungal hyphae underlying its potential mode of action. However, additional validation through large-scale and multi-environment field trials is required to confirm its efficacy, environmental safety, and long-term contribution to sustainable crop protection strategies.

## Acknowledgements

We would like to acknowledge CropBiome and UCD Conway Metabolomic Facility for their support. CropBiome have an exclusive evaluation and option licence agreement for 2Py-5CA. Australian patent application filed, 2024227203 and UK patent application GB 2414807.4. “ An antifungal composition and its uses”.

## Credit authorship contribution statement

Seema Rathore, Angela Feechan, Olga Lavostveky: Designed experiments. SR, OL, SJK and Junhao Xie performed the experiments. SR and OL carried out analysis of the results. AF and SR wrote and revised the manuscript. All authors contributed to the article and approved the submitted version.

## Supplementary Figures Legends

**Fig. S1.** Previously collected 64 endophytes^33,34^ were screened for inhibition of *Ramularia collo-cygni (RCC)* growth *in vitro* (A); Inhibition zone of *R. collo-cygni* around Ef20, Ef30 Ef65 and Ef39 The image showed Asterisks representing agar plugs of endophytes. (B) Percent (%) surface area covered by the growth of *R. collo-cygni* in the presence of endophytic fungi (Ef20, Ef30, Ef65 and Ef39 (negative control) measured using ImageJ (https://imagej.net). Mean ± SD; different letters represent statistically significant differences One-Way ANOVA Tukey’s tests (*P* ≤ 0.05) (C ).

**Fig. S2.** Fungal growth of the isolated endophytic fungal strains Ef20(A), Ef30 (B) and Ef65 (C). Colony cultures on potato dextrose agar incubated at 28 °C for 7 days. Light micrograph showing single conidiophores, stained with lactophenol cotton blue (D). Scale bar=100 μm.

**Fig. S3.** Interaction pattern between fungal endophytic fungi with *R. collo-cygni (RCC)*, *P. teres (Pt), F. graminearum (Fg)* on PDA (A-L). Light micrographs show smooth hyphal growth (black full arrow (A-C), new hyphal buds (dashed arrow, (A)) in the pathogen only (*R. collo-cygni, P. teres and F. graminearum)* panels (left). Hyphae of all the fungal pathogens (*R. collo-cygni, P. teres and F. graminearum)* are shrunken, vacuolated (arrowhead only), distorted and conidia are intertwined on the hyphae (small arrowhead, (D-L) treated with endophytic fungal isolates. Scale bar = 100µm.

**Fig. S4.** Antifungal activity of culture filtrate (CF) of endophytic fungal isolates *P. oxalicum* Ef20, Ef30 and *P. virgatum* Ef65 against *R. collo-cygni (RCC). RCC* growth on PDA control (A) CF of Ef30 showed strong inhibition of *RCC* colonies (B) CF of Ef20 with moderate inhibition and (C) CF of Ef65 with no inhibitory effect (D).

**Fig. S5**. Biocontrol efficacy of *P. oxalicum* Ef30 aqueous extract against *Ramularia collo cygni (RCC)* in dual plate disc assay. Inhibitory effect (clear zone) observed on the growth of *R. collo-cygni* by Ef30 aqueous extract on a paper disk compared to (A) control with *RCC* only and (B) *RCC* growth in the presence of the Ef30 ethyl acetate extract showing no activity on a paper disk (C).

**Fig. S6.** Graphic representation of treatments. Foliar spray of barley plants with *R. collo-*cygni (A); *RCC*+DMSO (0.5%); (B) RCC + different dosage of 2Py-5CA from 10 µg/mL – 80 µg/mL ; (C) pure antifungal compound 2Py-5CA alone (D) and only DMSO (0.5%) as negative control with no fungal pathogen (*RCC*) (E). 2Py-5CA was sprayed after 48h of post-inoculation of plants with RCC.

**Figure S7.** KEGG pathway mapping of Proline metabolism (https://kegg.jp/pathway/ko00330). Purple box-surrounded markers indicate genes involved in the biosynthetic pathway of L-Proline/2-pyrrolidone-*di*-carboxylic acid. The fluorescent green highlighted box around the formation of 1-Pyrroline-5-carboxylate (1-Py-5-CA). The phqD gene encodes L-1-pyrroline-5-carboxylate reductase (P5CR), an enzyme that plays an important intermediate role in the L-proline biosynthetic pathway.

## Supplementary Tables

**Table S1**. Screening of 64 endophytic fungal isolates inhibiting *R. collo-cygni* colonies growth. The strongest candidates were identified based on inhibiting *RCC* colonies. Using the ImageJ software confirmed *Penicillium oxalicum* Ef20 and *P. oxalicum* Ef30 as the two most potent isolates exhibiting significant inhibition of *R. collo-cygni* colonies growth. (See Excel File of Inhibition Scores. Supporting Information for Publication File)

**Table S2.** List of 18 common compounds found in the aqueous fraction of P. oxalicum Ef20 and Ef30, identified through untargeted metabolomics using LC-MS/MS. The compound fragments match with the Human Metabolome Database (HMDB) and MELTING database.

**Table S3.** Aqueous fraction identified secondary metabolites from Ef20 and Ef30 tested against fungal pathogens. All compounds listed were tested against *R. collo-cygni.* Only 2-Pyrrolidone-5-carboxylic acid (2Py-5CA) showed a strong inhibitory effect against a broad range of fungal pathogens of crops (*R. collo-cygni*, *Z. tritici*, *F graminearum* and *Pyrenophora teres*) and fruit pathogen (*B. cinerea*).

**Table S4**: The number and type of secondary metabolite biosynthetic gene clusters (BGCs) in endophytic strains *P. oxalicum* Ef20, *P. oxalicum* Ef30 and *P. verruculosus* Ef39.

**Table S5:** Prediction of common six secondary metabolites biosynthetic Gene Cluster (BGCs) exhibited 80 - 100% sequence similarity identified in the genomes of *Penicillium oxalicum* EF20 and Ef30. The identified BGCs and their correspondence to observed metabolites in the MIBiG database.

