## Supplementary figures and images for "Biocontrol efficacy of 2-pyrroldione-5-carboxylic acid (2Py-5CA), an antifungal bioactive macromolecule from the endophyte *Penicillium oxalicum*, against *Ramularia collo-cygni* in barley"

### Figure_ S2.tiff

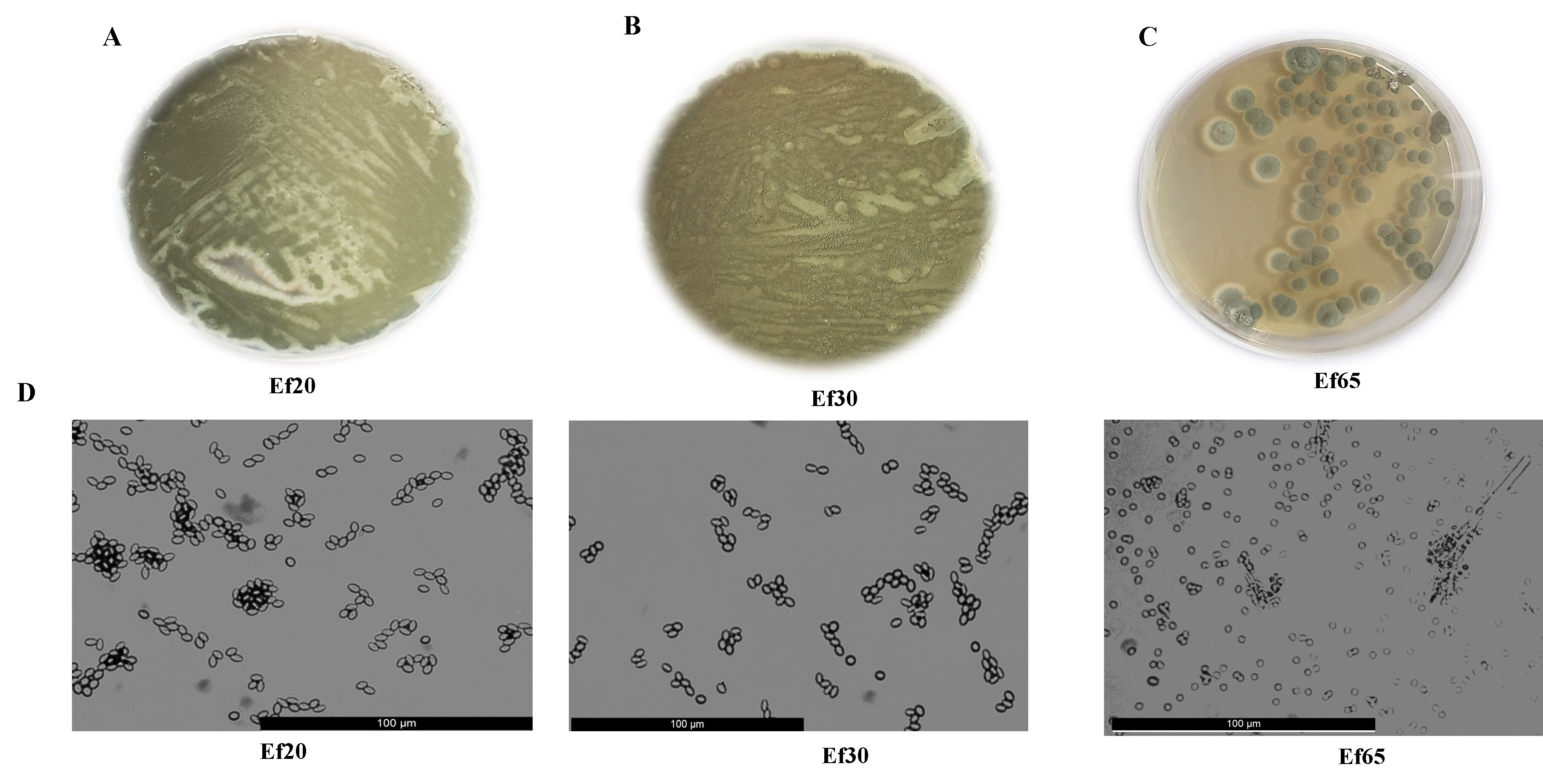

### Figure_ S5 .tiff

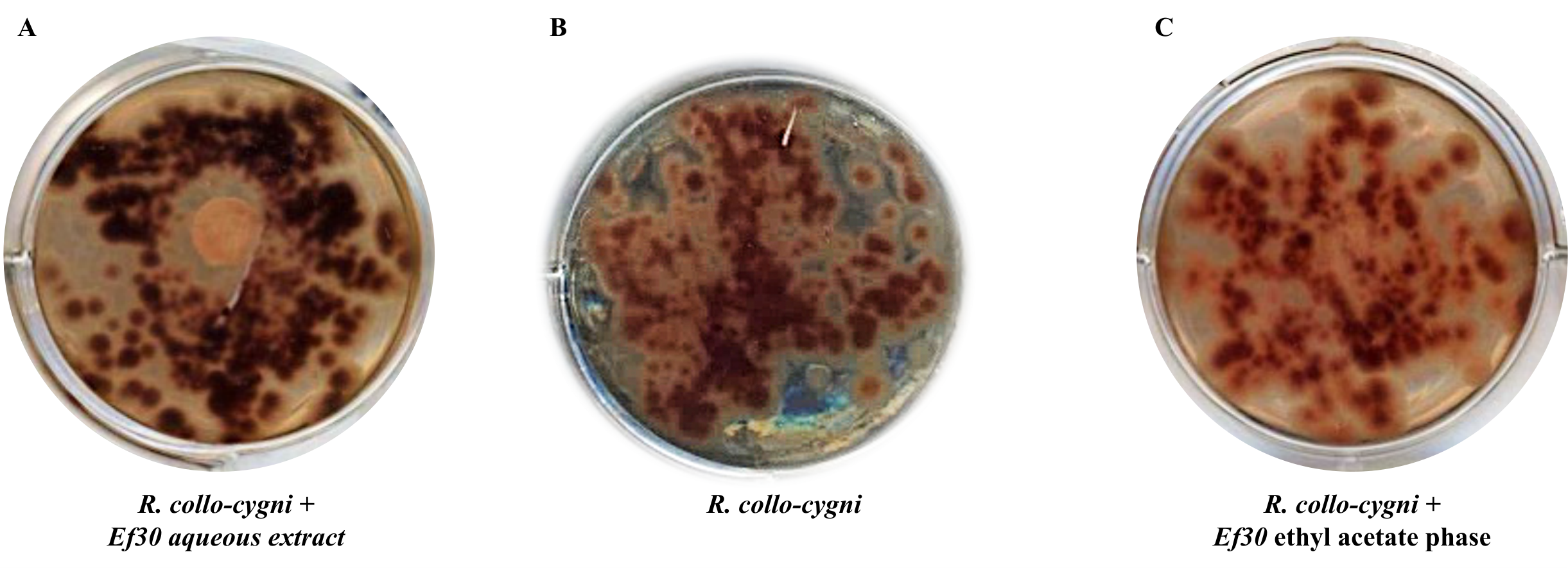

### Figure_+S1.tiff

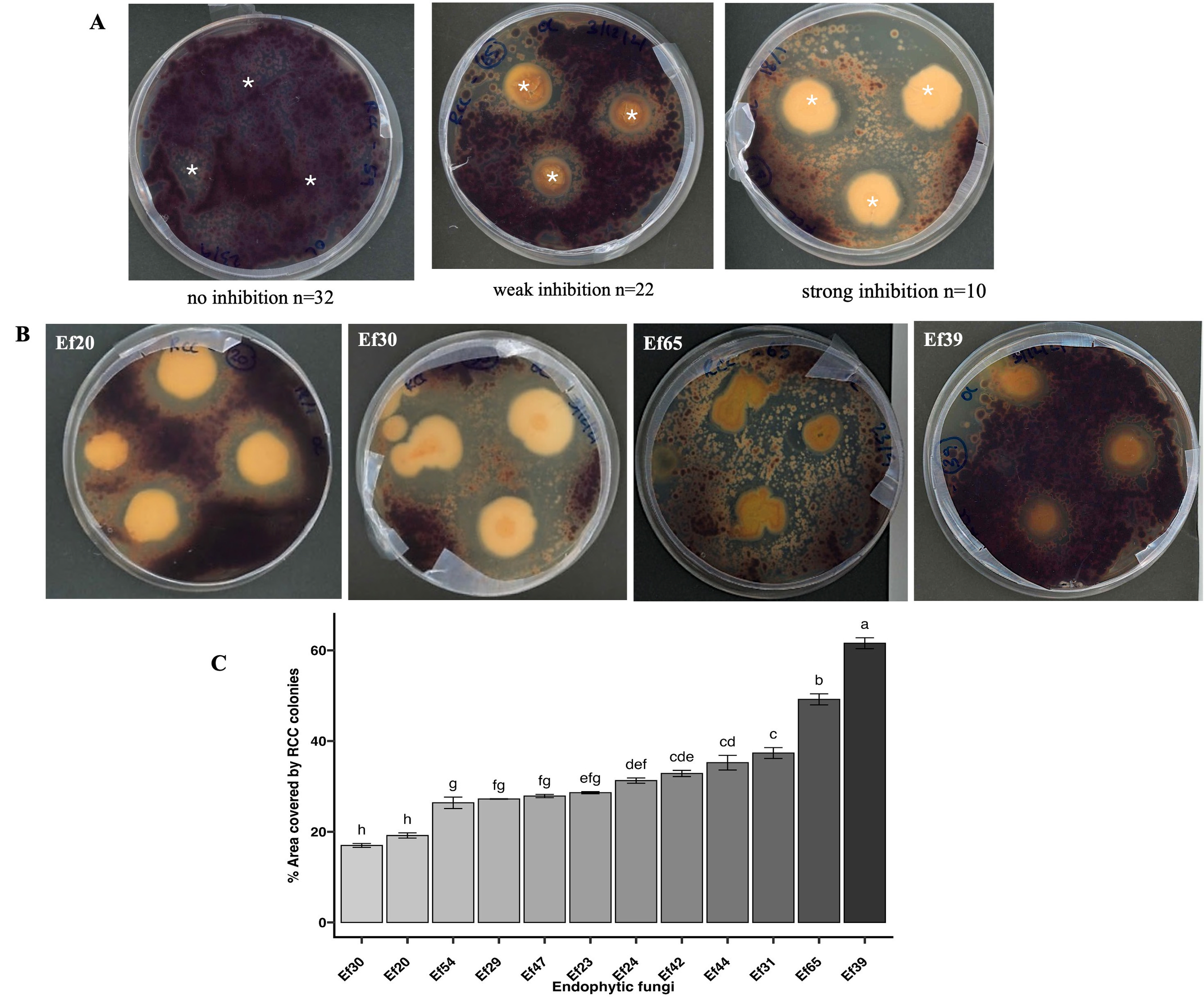

### Figure_+S3.tiff

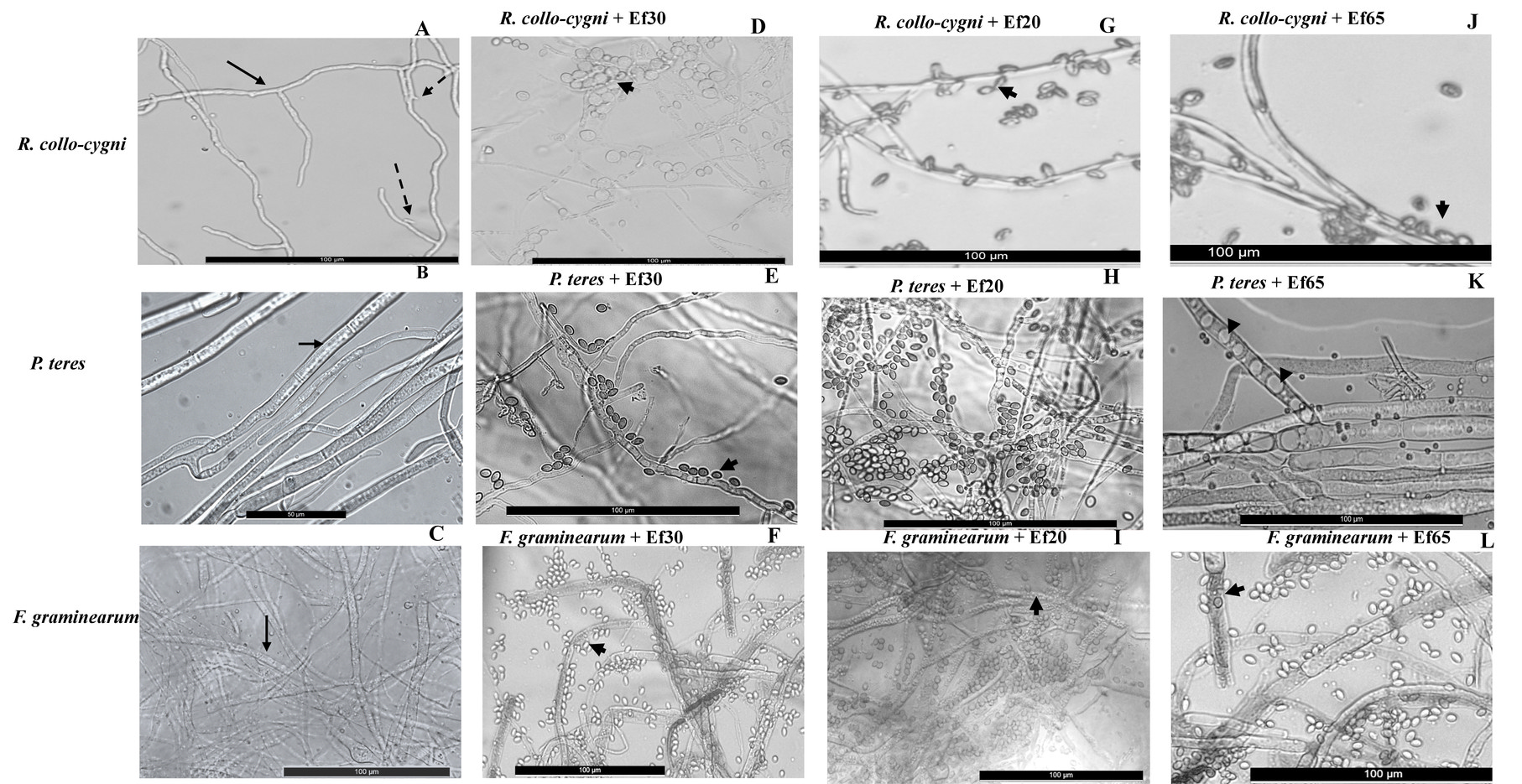

### Figure_+S4.tiff

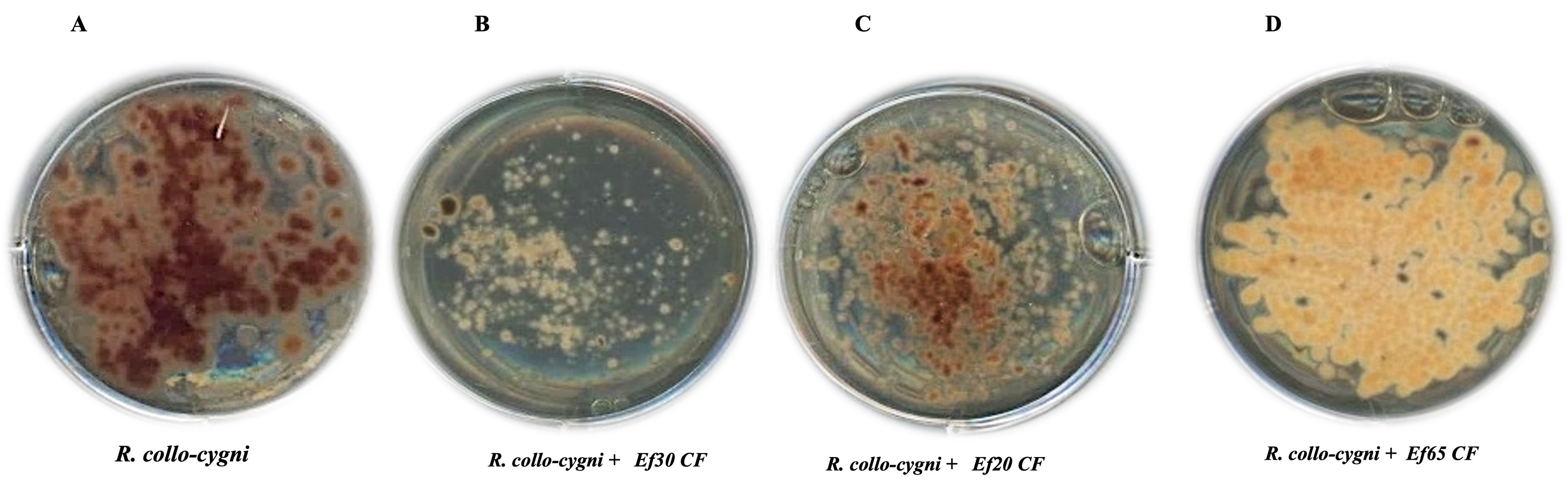

### Figure_+S6.tiff

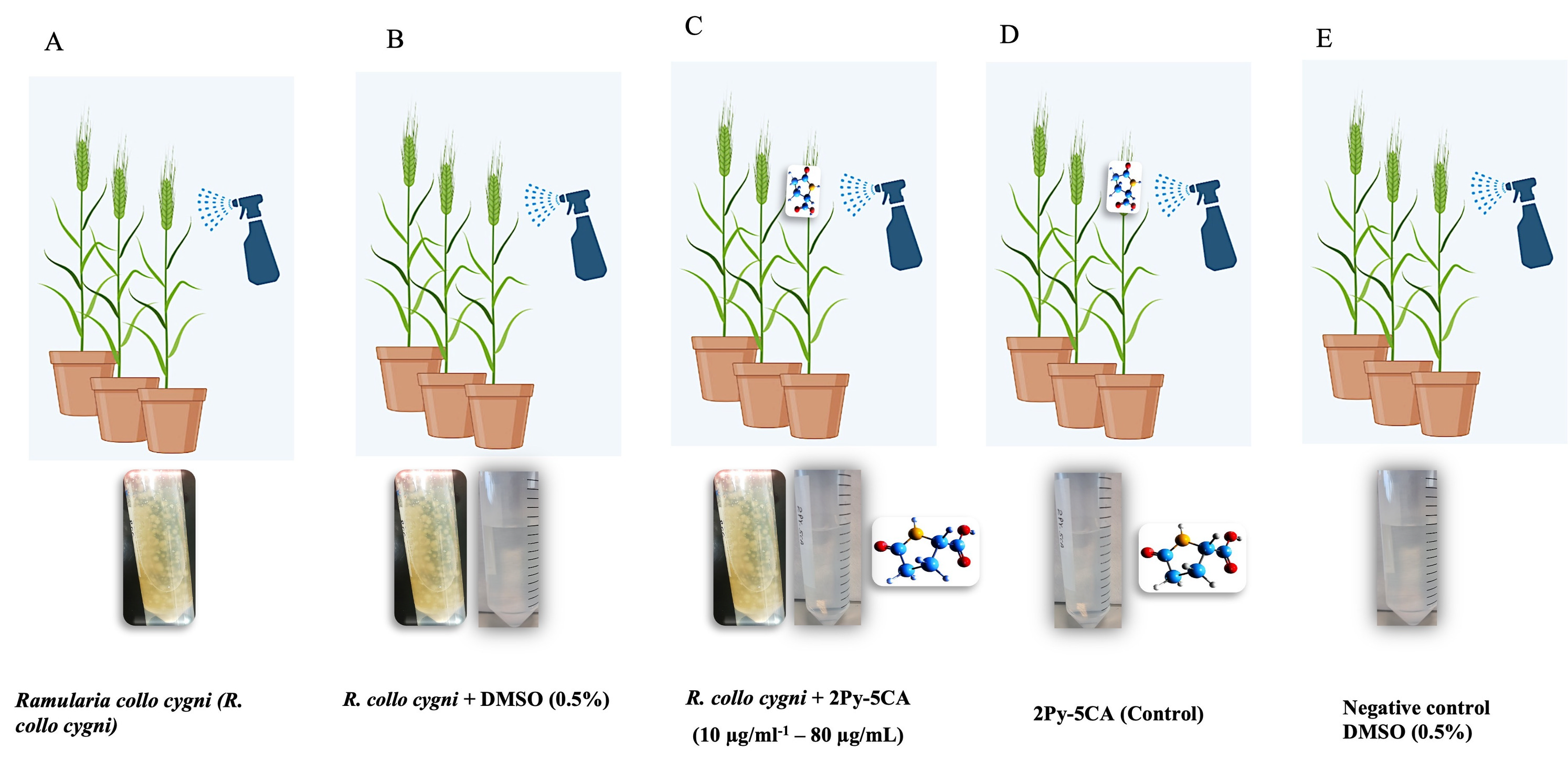

### Figure_+S7.tiff

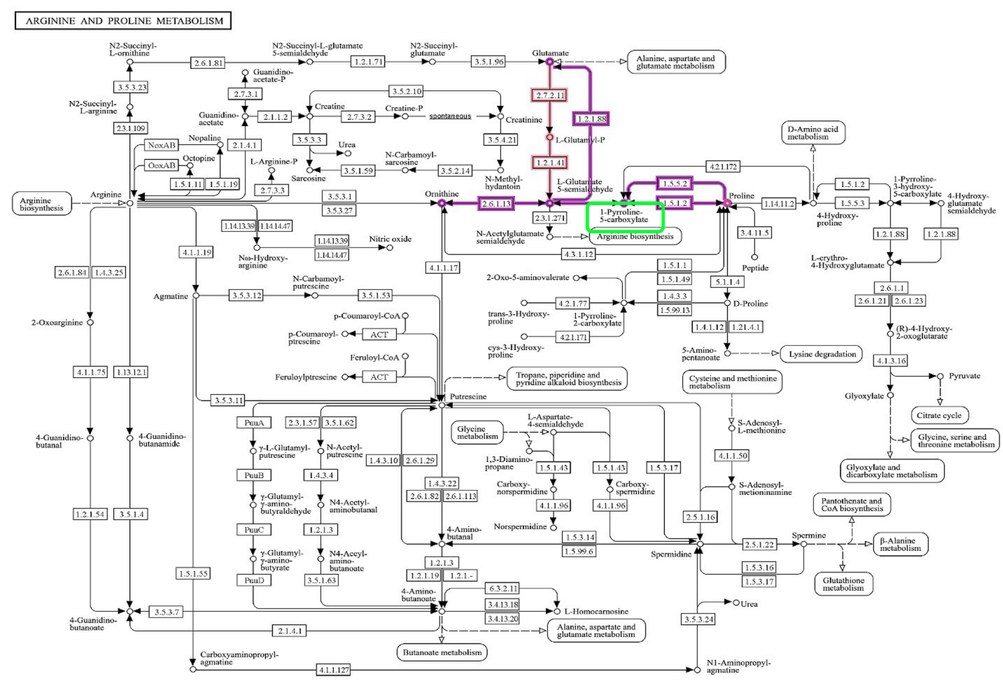

### Supplemental Table S2

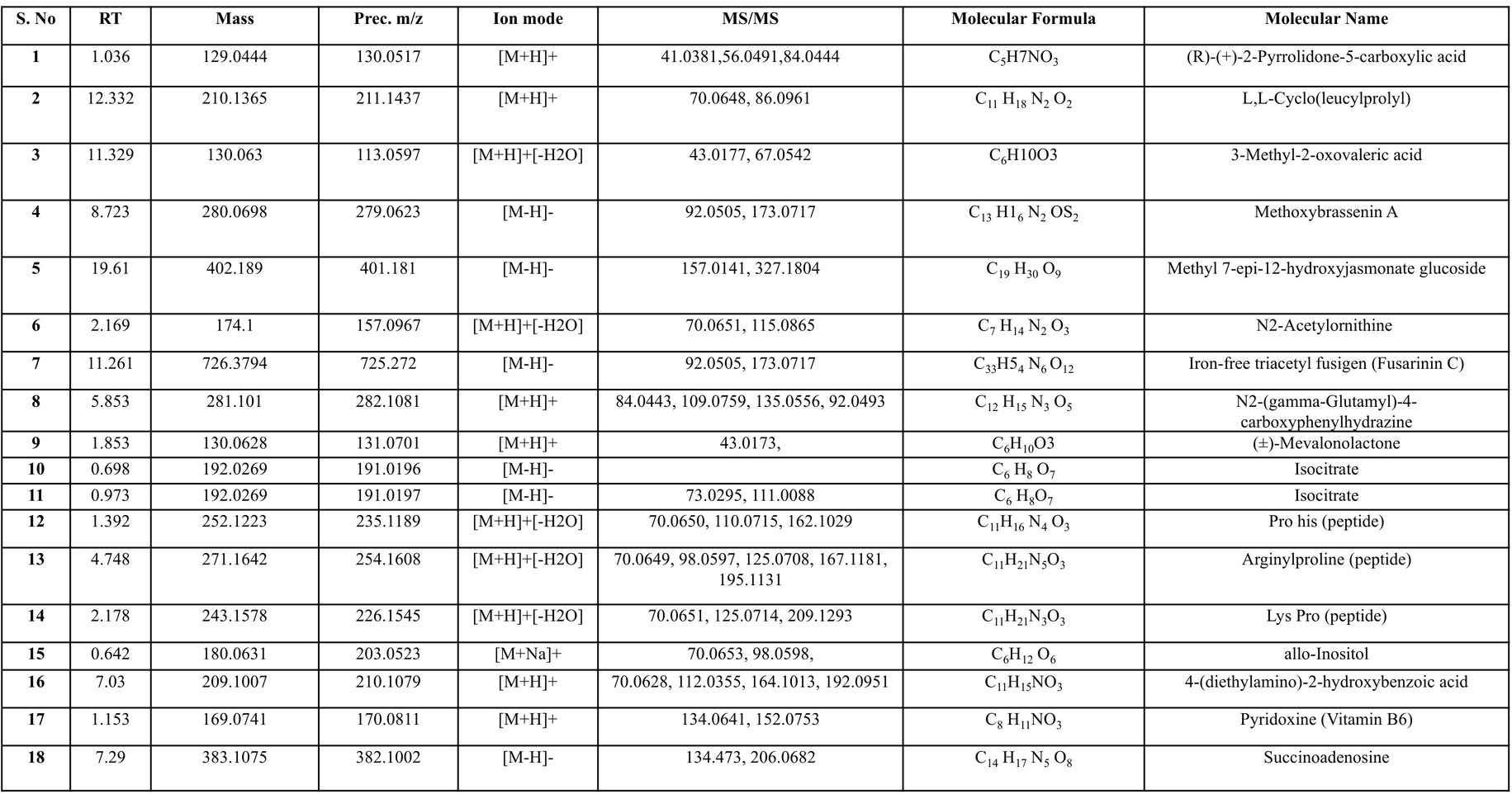

### Supplemental Table S3

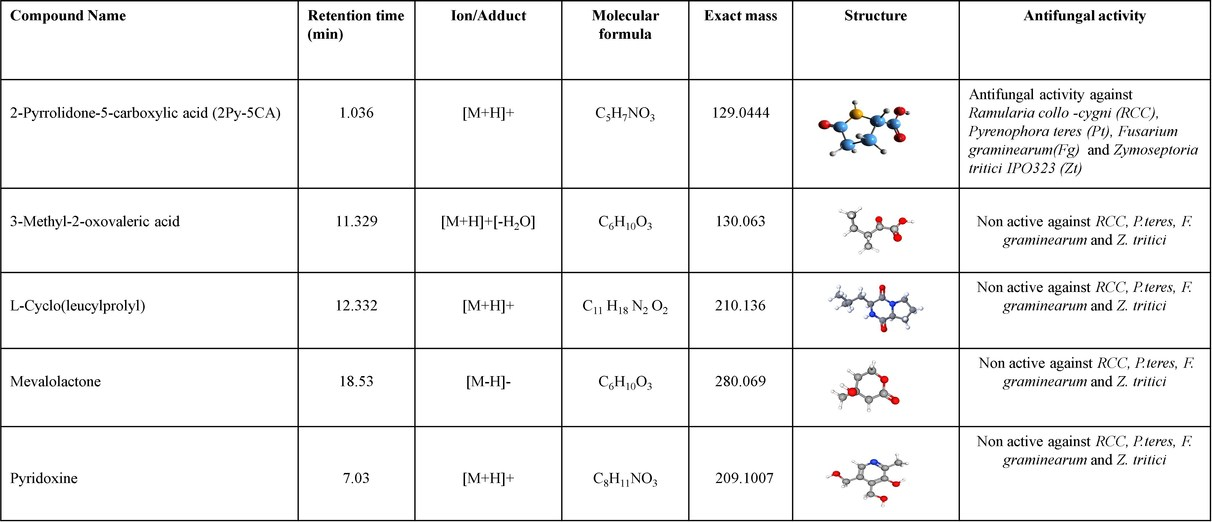

### Supplemental Table S4

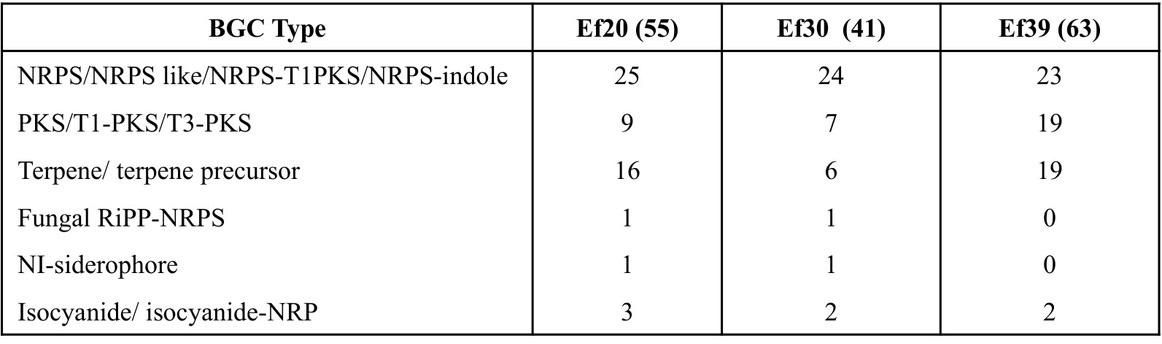

### Supplemental Table S5

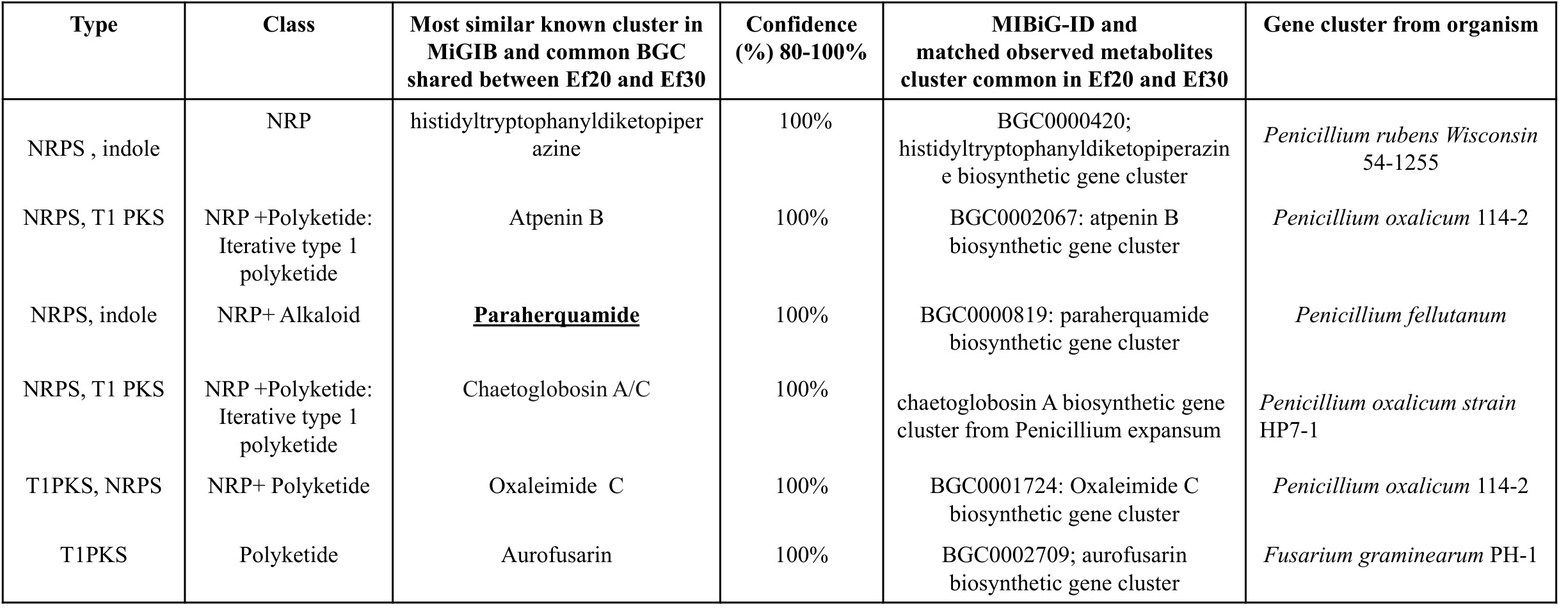
